# Age-associated aberrations of cumulus-oocyte interaction and microfilamentous structure in the zona pellucida decline female fertility

**DOI:** 10.1101/2023.12.30.573680

**Authors:** Yu Ishikawa-Yamauchi, Chihiro Emori, Hideto Mori, Tsutomu Endo, Kiyonori Kobayashi, Yuji Watanabe, Hiroshi Sagara, Takeshi Nagata, Daisuke Motooka, Akinori Ninomiya, Manabu Ozawa, Masahito Ikawa

## Abstract

One of the major age-related declines in female reproductive function is the reduced quantity and quality of oocytes. Here we demonstrate that structural changes in the zona pellucida (ZP) decrease fertilization rates from 34- to 38-week-old female mice, equivalent to the mid-reproductive of human females. In middle-age ovaries, the decline in the number of transzonal projections was accompanied by a decrease in cumulus cell-oocyte interactions, resulting in a deterioration of the oocyte quality. Scanning electron microscopy showed the ZP surface microfilament structure transitioning from rugged to smooth with age, leading to decreased fertilization rates due to impaired sperm binding to the ZP. Moreover, the fertilization rate of middle-age mice was restored to a comparable level to that of young mice by destabilizing the ZP in the presence of glutathione. These results suggest that the age-related structural changes in the ZP is a key for successful fertilization at reproductive age.

## Introduction

Aging is a complex biological process accompanied by physical changes throughout a lifetime. The functional decline of cells and tissues begins earlier in the female reproductive organs with individual aging (Broekmans et al., 2009; Heffner, 2004). One of the age-specific declines in tissue function, menopause, which is the loss of follicular function in the ovaries, typically occurs around age 50. In contrast, a marked decline in fertility is evident after age 35, with very limited pregnancies after age 40. The fertility rate for non-contraceptive populations is approximately 40% in the 20s to early 30s, 20-30% in the mid-30s, and 5-10% after age 40 among women (Heffner, 2004; Michelle J.K. Osterman, 2022).

In mammals, females undergo oocyte proliferation in the ovary during fetal development, and the oocytes are then stored in the ovary until sexual maturity and ovulation (Macklon & Fauser, 1999). The oocytes remain arrested in the meiotic first division within the primary follicle (McGee & Hsueh, 2000). As only a limited number of oocytes are activated and developed for ovulation, many follicles in the ovary remain dormant for years to even decades (Macklon & Fauser, 1999). Recent comprehensive analyses using RNA-seq have unveiled gene expression patterns in oocytes correlated with aging in mice and humans (Babayev & Duncan, 2022; Hamatani et al., 2004; Llonch et al., 2021; Pan et al., 2008). Indeed, DNA repair disorder, telomere shrinkage, chromosome missegregation, and mitochondrial dysfunction are predominantly involved in the deterioration of oocyte quality in mice aged 12-20 months (late-reproductive age, equivalent to age 40-60 in humans) (Dutta & Sengupta, 2016). While correlations between aging and the quality of oocyte have been found in late-reproductive age, why fertility begins to decline from mid-reproductive age remains unclear. A previous study demonstrated that the number of offspring in spontaneous pregnancies decreases after 8 months in mice (mid-reproductive age equivalent to age 35-40 in humans) (Suzuki et al., 1994). The cumulus cells also showed age-specific gene expression changes earlier than the oocyte with aging (Mishina et al., 2021), presuming that the maintenance of functional cumulus cells is critical to assuring oocyte quality.

During late folliculogenesis, the oocyte has physical contact with granulosa cells (progenitor of cumulus cells) and forms the cumulus-oocyte complex (COC) at ovulation. Their close interaction with the oocyte during folliculogenesis contributes to the maturation of the oocyte. Cumulus cells extend multiple filopodia into the zona pellucida (ZP) in the oocyte, known as a transzonal projection (TZP), making bridges between cumulus cells and the oocytes. Cumulus cells play a role in cellular communication, including endocrine, autocrine, and paracrine regulators such as amino acids and ions, which facilitate follicle development and ovulation (Eppig et al., 2005). Its failure interferes with oocyte maturation and fertility (Kidder & Mhawi, 2002; Simon et al., 1997). However, little is known about how aging of cumulus cells is involved in the maturation of the oocyte.

As mentioned above, a gap between individual aging and reproductive aging is highly recognized, understanding of fertility decline in middle-aged females is essential to elucidate the mechanism of reproductive aging. In the present study, we demonstrated how aging affects the intercommunication between cumulus cells and the oocyte and subsequent fertilization using middle-aged (34- to 38-week-old) female mice. We found that, in the ovary, the oocyte quality declines due to a lack of interaction between the cumulus cells and the oocyte. In aged oocytes, the lack of ZP surface mesh structure prevents sperm binding and penetration. Lastly, adding glutathione to loosen ZP rescued fertilization rates. These results suggest that age-related oocyte-cumulus miscommunication impairs the formation of ZP and middle-aged female fertility.

## Results

### Middle-aged female mice show a reduction of ovarian reserve of follicles and ovulated oocytes

When we examined the estrous cycle by taking daily vaginal smears, diestrus was prolonged in middle-aged female (middle) mice compared with young-aged female (young) mice (The average number of estrus cycles was 3.2±0.2 and 2.3±0.1 during a 20 day-test in young and middle, respectively), still the estrus cycles were maintained (Figure S1A-D). We next investigated female fecundity by natural mating. We found that the number of pups in middle-aged mice was significantly reduced compared to that in young mice despite successful coitus demonstrated by vaginal plug formation (Figure 1A). Morphological analysis revealed that ovarian tissues of middle mice turned brown, while the oviduct and uterus and the ovarian weight remained comparable between young and middle-aged mice (Figure 1B and C). However, the area of ovarian fibrosis (Briley et al., 2016; Foley et al., 2021) was highly abundant in middle-aged mice (Figure 1D and E). Furthermore, the number of follicles in middle-aged mice was significantly decreased among primordial, primary, and secondary follicles (Figure 1F and G). When we retrieved the oocytes from oviducts after superovulation in middle-aged mice, half as many oocytes as in young mice were recovered (Figure 1H and S1E). Because the hormone levels in the sera of young and middle-aged mice were comparable in estradiol (Figure 1F), we focused on the development of antral follicles. As a result, the number of antral follicles (follicular diameter > 300 µm) was significantly reduced 48 hours after pregnant mare’s serum gonadotropin (PMSG) treatment in middle mice (Figure 1J and I). These results indicate that aging at the middle reproductive stage impairs follicular development in the ovary.

**Figure 1.**
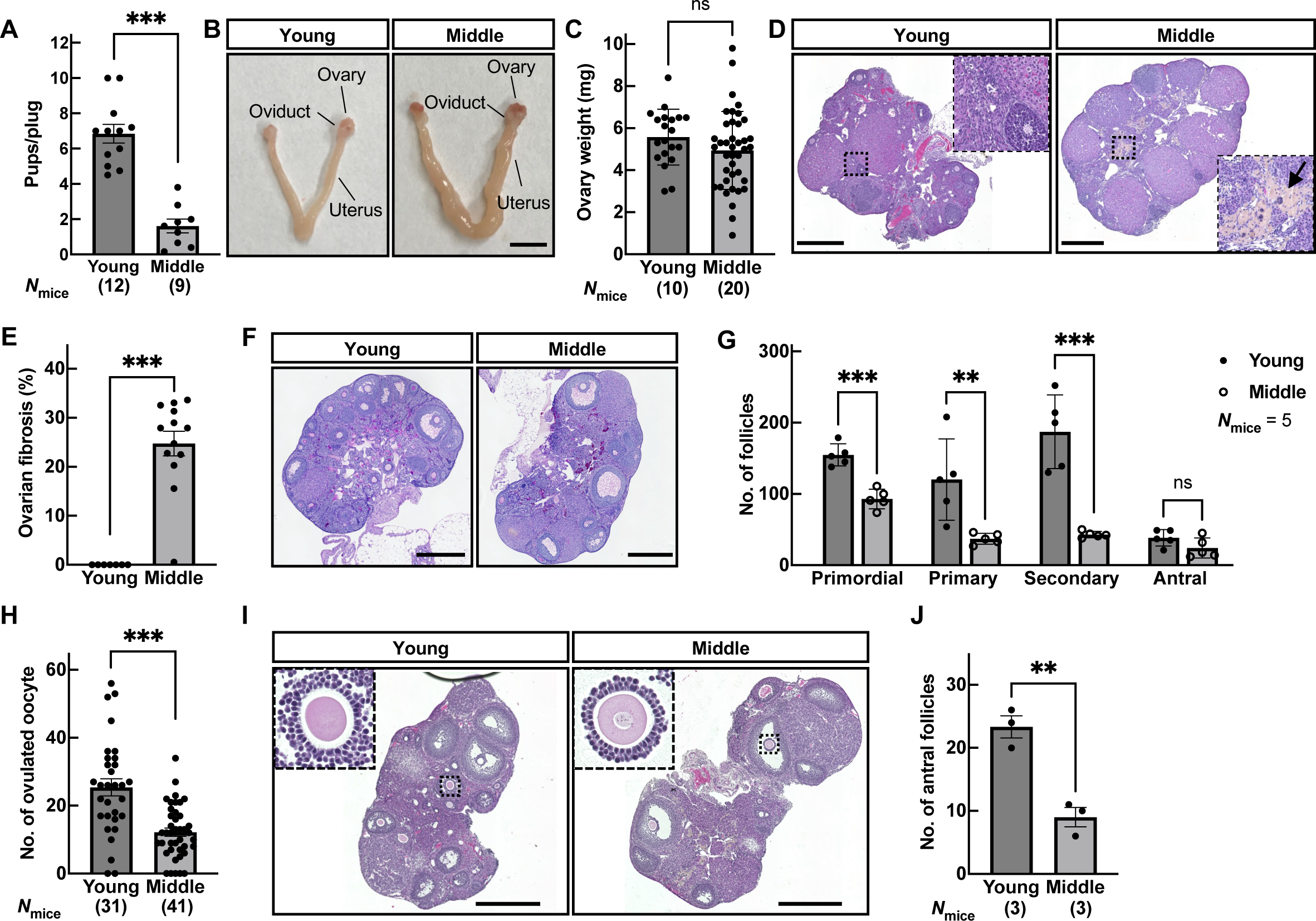
Physiological aging reduced ovarian reserve of follicles and ovulated oocytes in middle-aged mice. (A) The fecundity of young-aged (10- to 12-week-old; young) and middle-aged (34- to 38-week-old; middle) female mice was analyzed by natural mating; n indicates the number of female mice examined. \*\*\**p* < 0.001; Tukey-Kramer test. (B) Tissue morphology of the ovary, oviduct, and uterus. Scale bar = 5 mm. (C) Ovary weight. ns, not significant (*p* > 0.05; Tukey-Kramer test). (D) Representative hematoxylin and eosin (HE) stained ovarian sections. The regions surrounded by dotted lines are highlighted. Arrow indicates the area of ovarian fibrosis. Scale bar = 500 µm. (E) The ratio of fibrosis area in the ovary. \*\*\**p* < 0.001; Tukey-Kramer test. (F) Representative hematoxylin – PAS stained ovarian sections. Scale bar = 500 µm. (G) The number of follicles including primordial, primary, secondary, and antral, in the ovary. \*\**p* < 0.01; \*\*\**p* < 0.001; Unpaired Student’s t-test. (H) The number of ovulated oocytes. Superovulation was induced with 7.5 IU (international units) each of PMSG and hCG at 48-hour intervals; n shows the number of females analyzed. \*\*\**p* < 0.001; Tukey-Kramer test. (I) Histological analysis of the ovary 48 h after PMSG injection. The regions surrounded by dotted lines, the cumulus-oocyte complex (COC), in young and middle ovarian follicles are highlighted. Scale bar = 500 µm. (J) Total number of antral follicles (max follicular diameter > 300 µm) from ovaries 48 h after PMSG injection. \*\**p* < 0.01; Unpaired Student’s t-test. Data are mean±SEM.

### Impaired intercommunication between cumulus cells and oocytes causes reduced oocyte quality during aging

The cumulus cells, somatic cells that surround the oocytes within the follicle, play an essential role in promoting oocyte maturations by bidirectionally transporting ions, amino acids, small molecules, and hormonal signaling through gap junctions during folliculogenesis (Matzuk et al., 2002). Previous studies demonstrated that cumulus cells undergo transcriptome changes during aging at the middle reproductive stage (Mishina et al., 2021) and secrete factors that deteriorate the oocytes in vivo and in vitro (Babayev & Duncan, 2022). We thus asked whether the bidirectional communication between the cumulus cells and oocytes was attenuated by aging in the follicle. When we observed the antral follicles using a transmission electron microscope (TEM), the cumulus cells appeared regularly arranged along the oocyte in young (Figure. 2A). In contrast, the density of cell adhesion between the cumulus cells was lost, and the spacing between the cumulus cells was significantly enlarged in middle-aged mice (Figure 2A, B, and closeup pictures).

**Figure 2.**
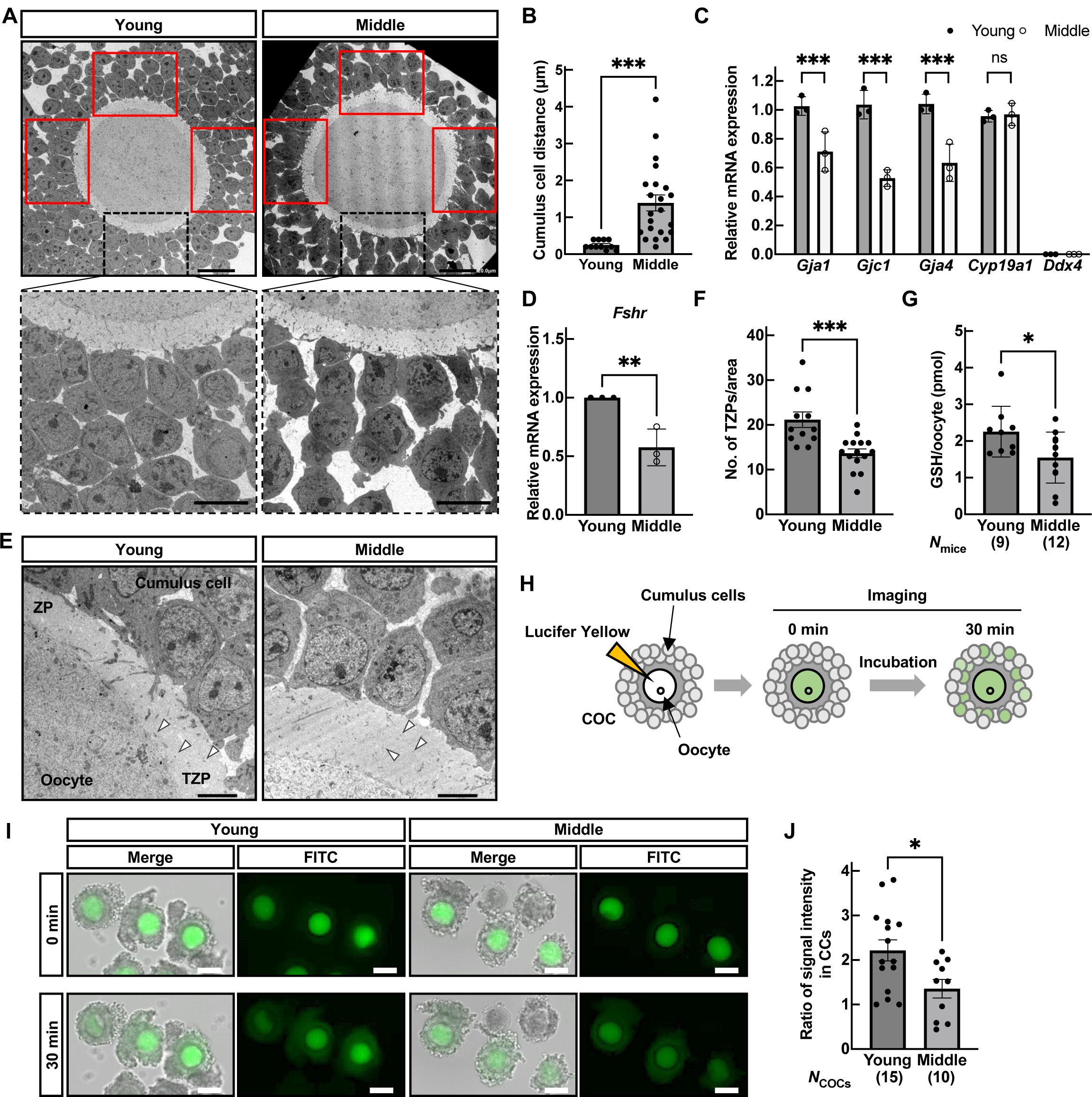
Aging leads to a depletion of cumulus cell gap junctions and the loss of intercommunication between cumulus cells and oocytes. (A) Representative TEM pictures of the antral follicles. The regions surrounded by dotted lines are enlarged at the bottom. Scale bar = 20 µm (upper), 10 µm (bottom). (B) Cell-to-cell distance between cumulus cells in the antral follicles. Four different positions, including red lines and black dotted lines in Figure 2A, are used for analysis***p < 0.001, Tukey-Kramer test. (C and D) Gene expression of gap junction genes including *Gja1* (Connexin 43), *Gjc1* (Connexin 45), *Gja4* (Connexin 37), cumulus cell maker *Cyp19a1*, germ cell marker *Ddx4*, and follicle-stimulating hormone receptor *Fshr* in cumulus cells. Cumulus cells were collected from antral follicles 48h after PMSG injection. ***p < 0.001; ns, not significant (p > 0.05), unpaired Student’s t-test. (E) TEM pictures of the COCs in the antral follicle, including cumulus cells, the zona pellucida (ZP), transzonal projection (TZP), and oocyte. Scale bar = 5 µm. Arrowheads indicate TZPs. (F) The number of TZPs. ***p < 0.001; Tukey-Kramer test. (G) Amount of glutathione (GSH) in an oocyte; n indicates the number of female mice examined. *p < 0.05; Tukey-Kramer test. (H) Schematic diagram of lucifer yellow injection into the GV oocytes with cumulus cells. Images were taken at 0 and 30 min after injection. (I) Pictures of the COCs at 0 and 30 min after injection. Scale bars = 100 µm. (J) The ratio of fluorescence intensity of cumulus cells before/after incubation; n indicates the number of COCs tested. *p < 0.05; Tukey-Kramer test. Data are mean±SEM.

The cumulus cells form gap junctions; the innermost layer of cumulus cells surrounding the oocyte extends cytoplasmic projections/filopodia, transzonal projections (TZPs), to transmit the essential factors to the oocyte for its maturation. A previous study suggests that aging impacts cumulus cells first then oocytes because transcriptomes change in cumulus cells and the GV oocytes at 9-month-old and 14-month-old, respectively (Mishina et al., 2021). We thus analyzed the published datasets (GSE159281) to characterize the expression patterns of gap junction genes involved in cumulus cell adhesion. Gene expression of *Gjc1* (Connexin 45) was significantly decreased at 9 and 14 months of age (Figure S2). In addition to *Gjc1*, gene expression of the gap junction gene, *Gja1* (Connexin 43) and *Gja4* (Connexin 37), which are also mainly expressed in the cumulus cells and function in cell adhesion of the cumulus cells, were found to be significantly downregulated in antral follicles of middle compared to that of young by qPCR (Figure. 2C). Because estrus cycles were difficult to be determined by smear test (Figure S1C and D), we next treated females with PMSG. The expression level of *Fshr*, which is expressed in the cumulus cells and induces follicle development in response to follicle-stimulating hormone, was significantly decreased in middle-aged mice (Figure 2D), regardless of comparable endogenous secretion of estradiol at the estrus stage (Figure S1F).

TZPs bridge cumulus cells and oocytes to exchange amino acids and nutrients in maturing GV oocytes, making TZPs the histological marker for intercommunication between cumulus cells and oocytes. When we carefully observed the ZP, the number of TZPs decreased by about half in middle-aged mice (Figure 2E F). To examine the intercommunication between cumulus cells and GV oocytes, we performed two types of experiments. First, we measured the amount of cytoplasmic glutathione (GSH) in GV oocytes (Figure 2G). GSH is a tripeptide composed of glycine (Gly), L-cysteine (L-Cys), and L-glutamic acid (L-Glu) residues, defined as one of the indicators of cytoplasmic maturation in oocytes, and plays antioxidant roles in the oocytes (Tatemoto et al., 2000). Moreover, it has been reported that the cumulus cells transport Gly, L-Cys, and L-Glu amino acids via TZPs to regulate GSH synthesis and accumulation in the oocytes (Mori et al., 2000), suggesting that cumulus cells are responsible for the maintenance of cytoplasmic GSH concentrations in the oocytes. The cytoplasmic GSH concentration varied among GV oocytes in middle-aged mice, but the average was significantly reduced compared with young GV oocytes; 2.28 and 1.55 pmol/oocyte in young and middle, respectively. In the second experiment, we injected lucifer yellow into GV oocytes and observed its transfer into the cumulus cells. The transitional ratio in middle GV oocytes was only about half that of young GV oocytes (Figure 2H-J). These results suggest that the dysfunction of cumulus cells accompanied by the disturbed cumulus cell layered structure leads to poor cumulus cell-oocyte intercommunication and oocyte quality.

### Impaired sperm binding to ZP results in reduced fertilization rates in midde-aged mice

We hypothesized that oocytes with reduced intercommunication with cumulus cells during folliculogenesis would affect their subsequent capacity for fertilization and development in middle-aged mice. We further analyzed the transcriptomes of young and middle post-ovulatory MII oocytes and cumulus cells using bulk RNA-seq (Figures 3A and S3A). The MII oocytes and cumulus cells showed their characteristic gene expression patterns (Figure S3B). Consistently with previous study (Mishina et al., 2021), the MII oocytes in middle-aged mice had no age-specific gene clusters compared to young MII oocytes (Figure 3B). In contrast to pre-ovulatory cumulus cells (Mishina et al., 2021), no age-specific gene clusters were found in ovulated cumulus cells (Figure 3B). Gene Ontology (GO) analysis also failed to list age-specific GO terms in biological processes (Figure S3C).

**Figure 3.**
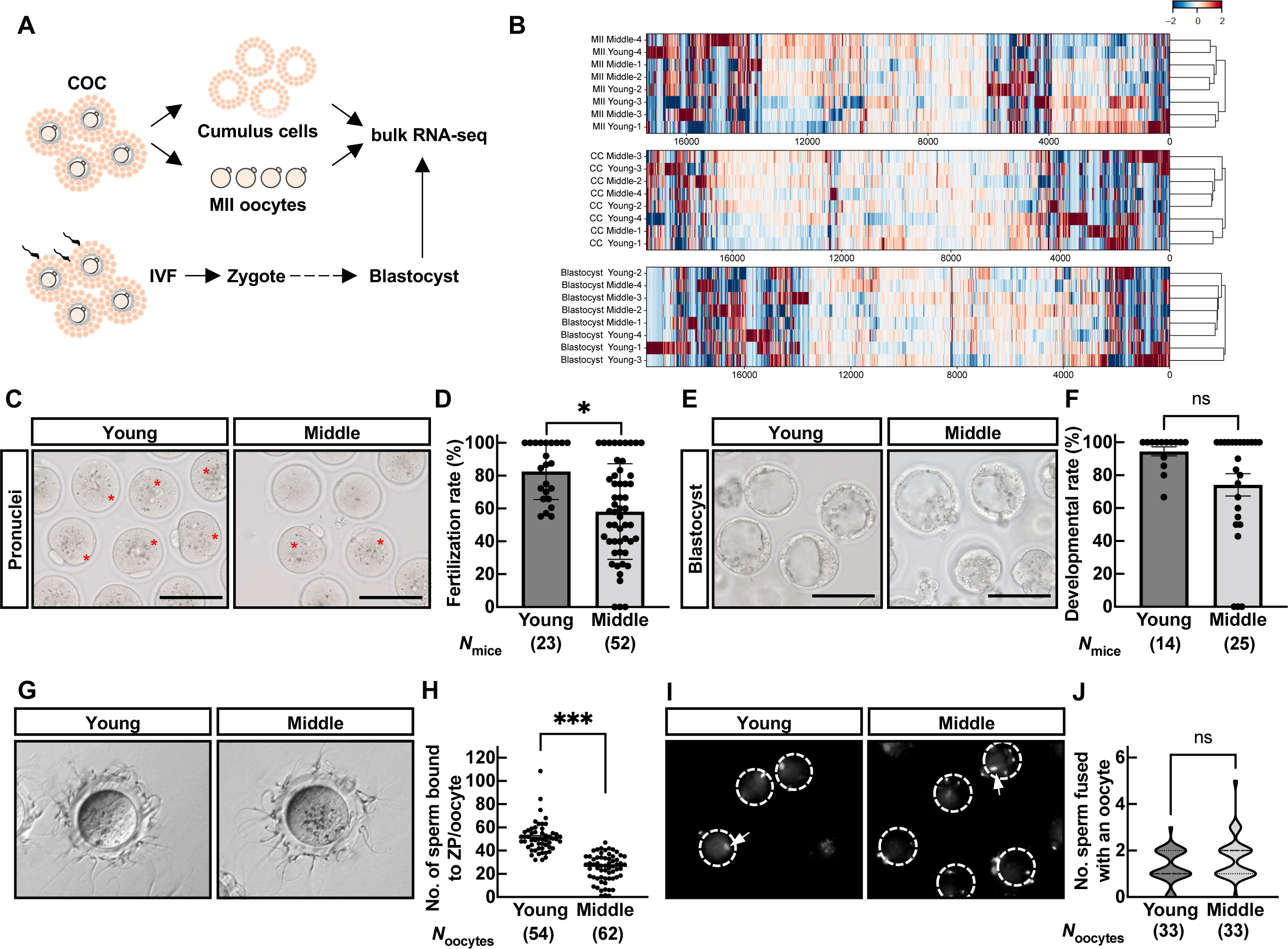
Fertilization rate decreased due to the failure of sperm binding to the ZP in mid oocytes. (A) Schematic diagram of bulk RNA-seq using MII oocytes, cumulus cells, and blastocysts. (B) Transcriptome analysis of MII oocytes, cumulus cells at ovulation, and blastocysts after IVF between young and middle mice. (C) IVF assay. Representative pictures of zygotes. Fertilized eggs were defined by pronuclei in zygotes. Asterisks indicate a zygote. (D) Fertilization rate. Fertilization rate was obtained after excluding fragmented oocytes. n indicates the number of female mice examined. \**p* < 0.05; Mann-Whitney test. (E) Representative pictures of blastocysts after 4-day incubation following IVF. Scale bar = 100 µm. (F) Developmental rate. Developmental rate was obtained by number of blastocysts over total 2-cell embryos. ns, not significant (*p* > 0.05; Mann-Whitney test). (G) Sperm-ZP binding assay. (H) Statistical analysis of the number of sperm bound to ZP. N indicates the number of oocytes used in each female. \*\*\**p* < 0.001; Tukey-Kramer test. (I) Sperm-oocyte fusion assay. Dotted circles and arrows indicate an oocyte and sperm nuclei fused with oolemma, respectively. (J) Statistical analysis of the number of sperm fused with an oocyte. ns, not significant (*p* > 0.05; unpaired Student’s t-test). Data are mean±SEM.

When we performed in vitro fertilization (IVF), the fertilization rate of mid MII oocytes was significantly lower than that of young (Figure 3C and D). When we cultured zygotes for four days, we found a tendency to decrease the developmental rate and a small difference in the transcriptome in middle-aged mice (Figures 3B, E, F, and S4A). However, there was no significant difference. To further elucidate which stage of the fertilization process is affected by aging (Figure S4B), we performed IVF with cumulus-free (sperm-ZP binding assay) or ZP-free (sperm-oolemma fusion assay) oocytes. Although an average of 50 sperm were bound to the ZP of young oocytes, the number of sperm bound to the ZP was decreased to half in middle MII oocytes (Figure 3G and H). Still, MII oocytes of middle-aged mice were fusion competent with young oocytes when freed from the ZP (Figure 3I and J). These results suggested that aging attenuates the fertilization potential of the oocyte by affecting the ZP rather than the oolemma.

### Aging alters the external microfilamentous structure of the ZP and affects fertilization

The ZP forms a mesh-like structure consisting of a delicate filamentous matrix and fibers. The ZP plays critical roles in fertilization, facilitating sperm binding, penetration, and preventing fertilization with extra sperm and heterospecific sperm. Since the oocytes are retained in the ovary until their activation for ovulation or atresia, we investigated how the structure of the ZP alters with aging. Taking advantage of scanning electron microscopy (SEM), we observed the ZP surface structure of MII oocytes immediately after ovulation and classified them into three groups (Figure. 4A). Type I; a well-defined reticular structure and fenestration with ruggedness on the surface. Type II; a reticular structure with tiny pores and less ruggedness. Type III; an entirely smooth surface with a collapsed mesh structure and no pores. Based on the classification, 95% of MII oocytes in young were classified as Type I, whereas the proportion of Type II and Type III MII oocytes was remarkably higher in middle-aged mice (Figure 4B). These results were consistent with a previous study (Nogues et al., 1988) and the presence of smooth structures in Type III was even more pronounced in MII oocytes of middle-aged mice.

**Figure 4.**
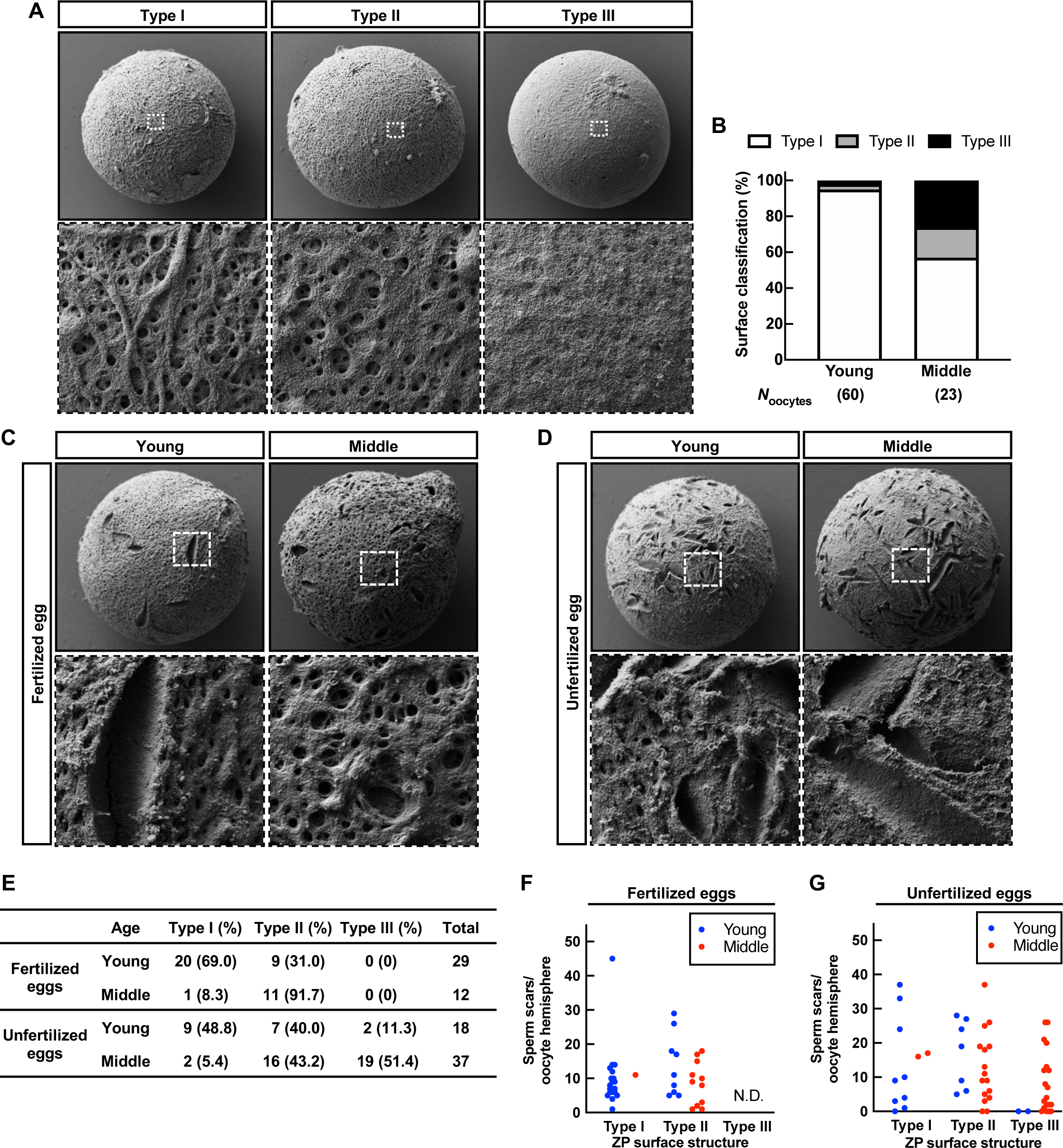
The ZP surface structure altered and affected fertilization with aging. (A) Classification of SEM images of ZP surface. The regions surrounded by dotted lines are highlighted at the bottom. Magnification x1500 in upper panels, and x10,000 in bottom panels. Type I surface structure has ruggedness with a network and spongy appearance containing numerous pores and fenestrations. Type II shows a more compact structure with fewer pores and smooth surfaces. Type III shows that fenestrations of the outer surface were rare or even absent. (B) The population of ZP surface classification based on the morphology of Figure 4A. (C and D) SEM images of ZP surface in fertilized eggs and unfertilized eggs. The regions surrounded by dotted lines are highlighted at the bottom. Magnification x1,500 in the upper panels, and x10,000 in the bottom panels. (E) The ZP surface classification of fertilized and unfertilized eggs. (F and G) The number of sperm tracks in the oocyte hemispheres.

Moreover, to objectively evaluate the ZP surface structure, we attempted to characterize the shape of the ZP surface by fractal dimensions, which represent universal quantities related to the geometrical or structural complexity. We developed an algorithm that analyzes surface ruggedness based on three validation factors: the fractal dimension (self-similarity or complexity), the number of holes that form between the mesh structures (a major characteristic of the ZP), and the edge intensity (unevenness) (Figure S5A). The algorithm was trained using leave-one-out cross-validation of each feature from the images and analyzed a total of 83 images. As a result, the algorithms achieved a classification accuracy of 87%. The likelihoods of the fractal dimensions were found to be 0.54 and 0.64 in young and middle, respectively (Figures S5B and C). The more rugged ZP surface structure of MII oocytes corresponded to a smaller fractal dimension and likelihoods, suggesting that young MII oocytes had a higher self-similarity and regularity. Conversely, the ZP surface structure became more complex with increasing likelihoods, with MII oocytes of middle-aged mice exhibiting more planar or smooth structures. These data indicate that structures subjectively classified as Type II and Type III showed a high fractal dimension and that the fractal dimension of the corresponding ZP surface structures could adequately explain the subjective observation. These results also suggest that aging increases the smoothing of the ZP surface structure of oocytes.

To investigate how alterations of the ZP surface structure affect fertilization, we retrospectively observed fertilized and unfertilized eggs, based on pronuclei formation after IVF, using a stereomicroscope and conducted SEM analyses. In young and middle-aged mice, most fertilized eggs were classified as Type I and Type II, respectively, with no eggs exhibiting Type III (Figure 4C and E), indicating that at least a reticular structure is necessary for fertilization. Unfertilized eggs showed fewer Type I oocytes and increased ratios of Type II and even Type III, regardless of age (Figure 4D and E). We also found some holes/scratches on the surface, so-called “sperm scars,” presumably undermined by sperm passing through the ZP (Yanagimachi, 1994). The number of sperm scars was approximately ten on the hemispheres of the fertilized eggs, and sperm numbers varied on the unfertilized eggs (Figure 4F and G). Since successful fertilization alters the ZP structure and prevents extra sperm entry (ZP block to polyspermy) (Fahrenkamp et al., 2020), the large number of sperm scars should represent the failure of fertilization. The findings suggest that one possible reason for the lower fertilization rate in IVF was that the surface structure of the ZP changes from ruggedness to smooth structure with age, and its formation prevents sperm binding to the ZP in middle-aged mice.

### Oocytes exhibit aberrant accumulation of ZP filaments in middle-aged mice

To investigate when the alteration of the ZP surface structure began to occur, we observed cross-sections of the ZP by TEM using GV oocytes in antral follicles. In the cross-sectional image of the ZP in young, the ZP fibers were disorderly arranged to extend (Figure 5A-C), as if diverging, toward the cumulus cells, and spaces not occupied by fibers were observed in the outermost layers of the ZP (Figure 5B, C, B’, and C’). The facts suggest that the ZP surface structure already has a mesh structure in GV oocytes in young and is maintained until fertilization. On the contrary, the surface of GV oocytes in middle-aged mice had no ruggedness (Figure 5D-F) because the fibers were densely packed and formed a flat line (Figure 5E, E’, F, and F’). Moreover, the gaps between fibers on the surface observed in GV oocytes in young were also absent (Figure 5E’ and F’). This result is consistent with Figure 4A that the surface structure observed in the MII oocytes did not exhibit a ruggedness structure in middle-aged mice. To objectively evaluate the degree of the ruggedness of the ZP surface in GV oocytes, we then analyzed the likelihoods using the algorithm developed in Figure 4. The scores showed that cross sections of the ZP in middle-aged mice were regular and less uneven; the average of likelihoods was 0.54 and 0.70 in young and middle, respectively (Figures 5G and S6A). The results indicate that even within follicular development, the ZP surface structure lost its ruggedness with age.

**Figure 5.**
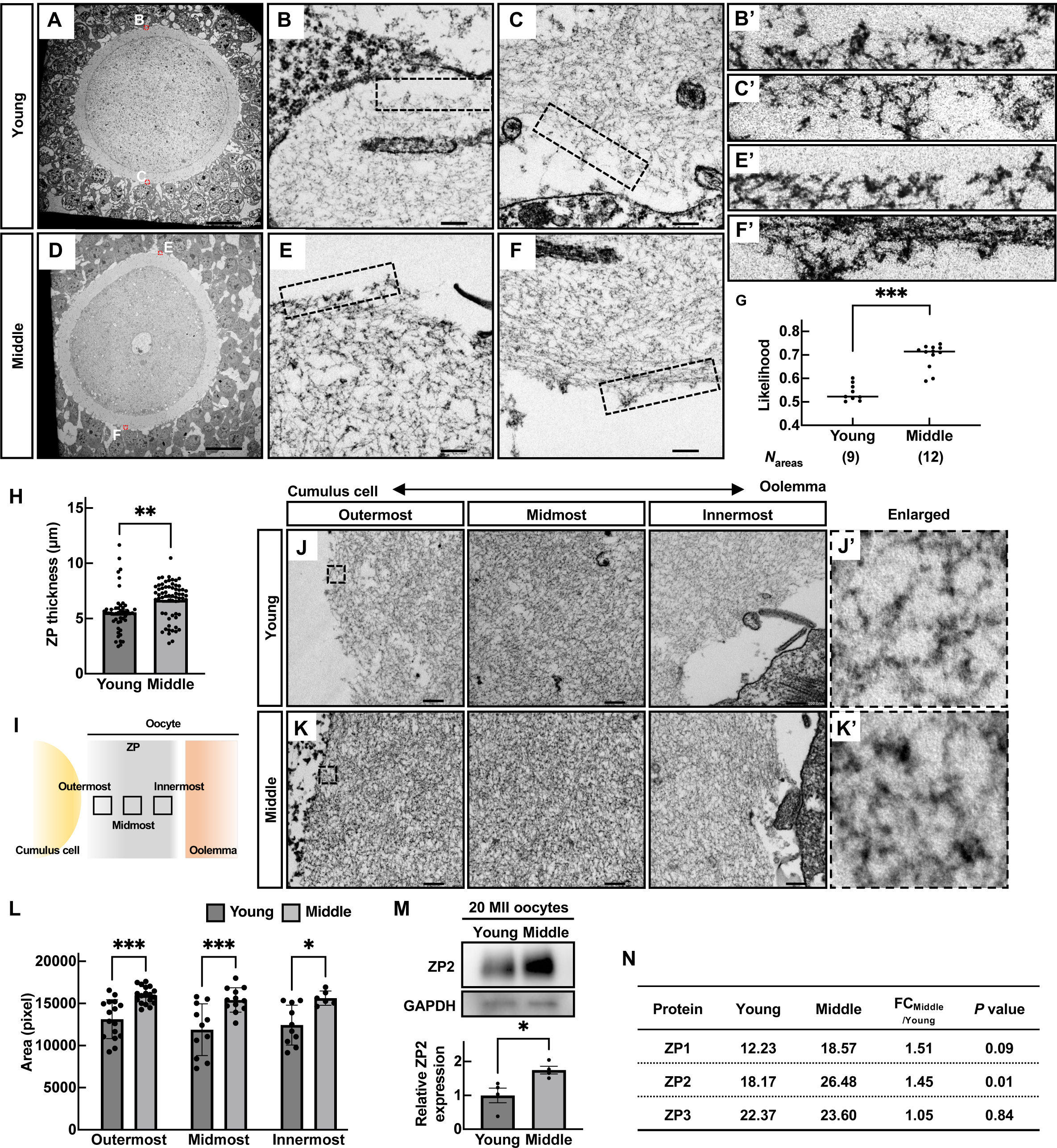
Aberrant accumulation of ZP filaments occurred in middle-aged mice. (A to F) TEM picture s of the antral follicles. The regions surrounded by red dotted lines are highlighted in B, C, E and F. Enlarged images of B, C, E, and F were also shown in B’ C’, E’, and F’. Scale bars = 200 nm in a and d, and 20 nm in B, C, E, and F. (G) Verification of surface roughness in the ZP. The value of likelihood was obtained by a fractal dimension analysis. (H) The ZP thickness. ** p<0.01; Tukey-Kramer test. (I) Schematic position of outermost, midmost, and innermost in the ZP. (J and K) TEM pictures of the ZP filaments composing the outermost, middle, and innermost areas. A double arrow explains a position between a cumulus cell side to an oolemma side. The regions surrounded by dotted lines are highlighted in j’ and k’. (L) Areas of ZP filaments in outermost, midmost, and innermost. *p< 0.05, ***p<0.001; Tukey-Kramer test. (M) Western blot of ZP2 in 20 MII oocytes. GAPDH was used as a loading control. Statistical analysis of ZP2. Four independent experiments were tested. *p< 0.05 (unpaired Student’s t-test). (N) MS analysis of the ZP. Average MS score was obtained from three independent experiments (*N*_mice_ = 3). p; unpaired Student’s t-test. Data are mean±SEM.

We further focused on the internal structure and protein expression of the ZP. Although there was no difference in the oocyte size and perivitelline space (PVS) in young and middle, thickness and total area of the ZP in MII oocytes were significantly increased in middle-age mice (Figures 5H and S6B-D). Comparing the density of ZP fibers in the assigned regions (outermost, midmost, and innermost), we found that the fibers were denser in all areas of the ZP in middle-aged mice (Figure 5I-L). These data indicate that the age-related change in the ZP affects the loss of surface roughness and the internal mesh structure.

The main components of the zona pellucida are ZP1, ZP2, and ZP3, with ZP2 being the most abundant component. We carried out western blotting and found that the ZP2 protein amount was 1.5 times greater in middle-aged mice than in young (Figure 5M). Given the unavailability of effective antibodies against ZP1 and ZP3, we further conducted MS analysis to compare the presence of those proteins quantitatively (Figure S6E). The results showed that the MS score of ZP2 increased in the ZP of middle-aged mice as in Figured 5M. In contrast, ZP3 and ZP1 showed an increasing tendency, but there were significant individual differences (Figures 5N and table S1). These indicate that aging causes changes in the surface structure and internal fiber occupancy of the ZP, at least with quantitative changes in the ZP constituent protein ZP2.

### Glutathione treatment restores the fertilization rate in middle-aged mice

Since the ZP structure changes with age, we tried to improve the fertilizing ability in middle-aged mice. Glutathione has been reported to reduce disulfide bonds of ZP proteins, increase free thiols, and thus destabilize the ZP, effectively improving the fertilization rate of frozen-thawed mouse sperm (Bath, 2010; Takeo et al., 2015; Takeo & Nakagata, 2011). When we incubated the COCs in a glutathione-containing medium, 98% of young MII oocytes showed ZP expansion, increasing the ZP diameter. The ZP diameter of MII oocytes also increased to an average of 125.3 µm, although there was some variation among the oocytes in middle-aged mice (Figure 6A and B). Moreover, as we carried out IVF with a glutathione-containing medium, average fertilization rates were recovered from 65.3% to 85.0% in middle-aged mice, comparable to young controls (Figure 6C and D). These results indicate that even though the surface structure and composition of the constituent proteins differ from young in aged oocytes, the loosening of the ZP could restore the fertilization rate in mice.

**Figure 6.**
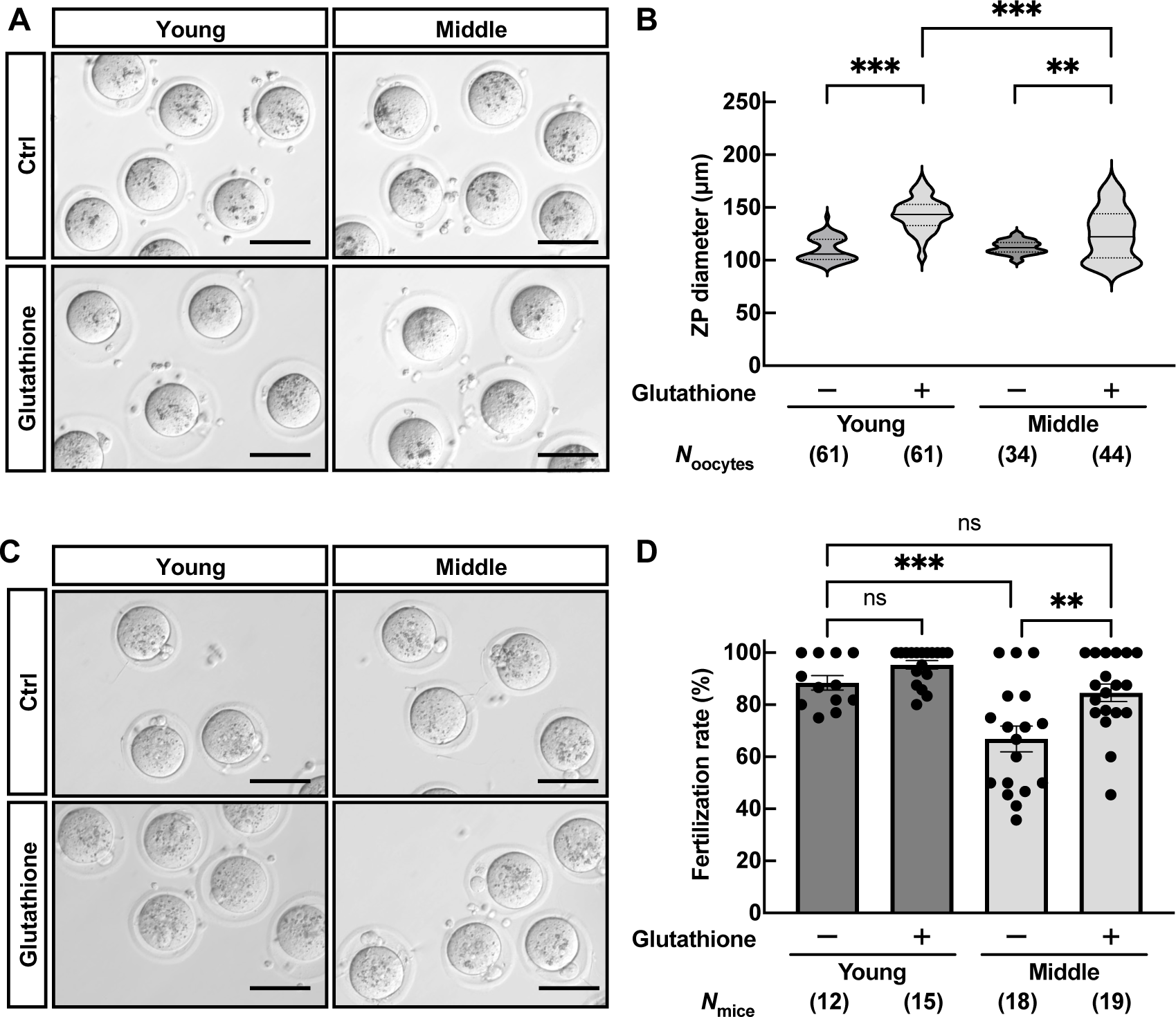
Glutathione treatment recovered the fertilization rate in middle-aged mice. (A) Representative pictures of the ZP expansion in the presence of glutathione. Scale bars = 100 µm. (B) ZP diameter; n indicates the number of oocytes examined. ** *p*<0.01; *** *p*<0.001; Tukey-Kramer test. (C) Representative pictures of fertilized eggs obtained through IVF in media containing glutathione. Scale bars = 100 µm. (D) Fertilization rate; n indicates the number of female mice examined. ** *p*<0.01; *** *p*<0.001; ns, not significant (*p*>0.05, Two-way ANOVA with Tukey’s multiple comparisons test). Data are mean±SEM.

## Discussion

Our present data show that an age-related reduction in oocyte-cumulus cell interactions in the ovary and subsequent structural changes in the ZP diminish fertilization at mid-reproductive age, which can partially be restored by glutathione treatment (Figure 7).

**Figure 7.**
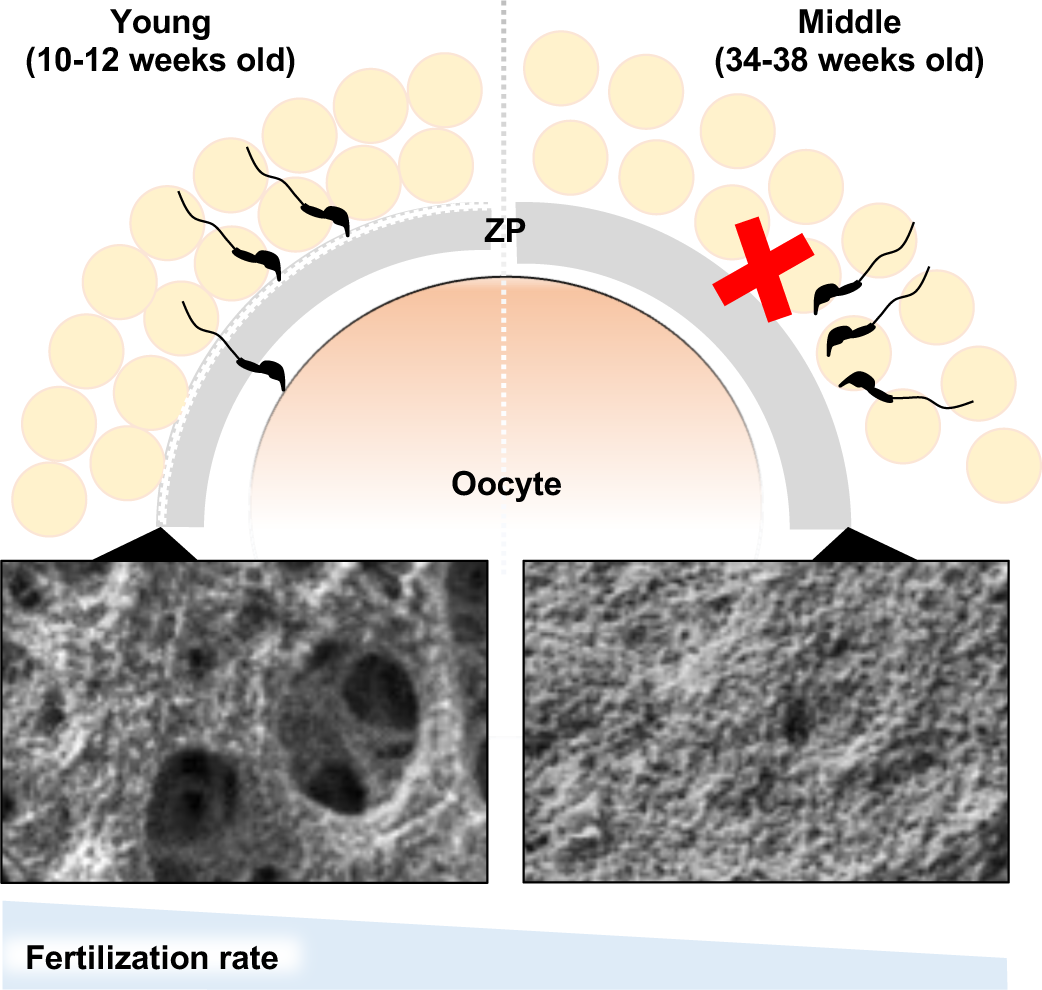
Schematic model of age-associated ZP structure affecting fertilization. In aging female mice, the quality of the oocytes was reduced in the ovary due to the decreased bidirectional communication between oocytes and cumulus cells. In addition, at least in mature follicles, the mesh structure of the ZP surface becomes a smooth structure, which persists during ovulation, preventing sperm from binding to the ZP and fertilization. The addition of glutathione in IVF could restore the fertilization rate of aged oocytes.

Previous studies have investigated the impacts of aging on female individuals, the ovaries, oocytes, and embryo development using naturally aged mice from 10 to 24 months of age (Bertoldo et al., 2020; Duncan et al., 2017). However, female mice fertility declines rapidly from 6 to 9 months (Suzuki et al., 1994), which is as early as six months before menopause (12-15 months) (Diaz Brinton, 2012; Russ et al., 2022). In the present study, our result reconfirmed that the fertility of middle-aged mice was significantly lower than that of young mice, despite their ability to produce pups. This result is consistent with humans; female fertility declines rapidly after age 35, while women reach menopause around age 50 (Broekmans et al., 2009).

As mentioned in Figure. 2, bidirectional communication between cumulus cells-oocytes had already diminished in the ovary during fertility decline. The COCs essentially form multilayered cumulus cells in the ovary; however, we found that these were thinner in middle-aged mice due to the weak adhesion between cumulus cells (Figures 1I and 2A). Intercellular communication is crucial for the oocyte maturation during folliculogenesis. Mutant mice lacking connexin 37 (Cx37, encoded by *Gja4*) exhibited infertile due to incomplete oocyte maturation at meiotic division by lack of gap junctions between cumulus cells and oocytes in follicles (Simon et al., 1997). Oocyte-specific deletion of connexin 43 (Cx43, encoded by *Gja1*) led to subfertility in mice (Gershon et al., 2008). The fate of premature oocytes goes to either atresia or failure of ovulation. Replacing Cx37-null mutant mice by specific expression of Cx43 in the oocytes restored the formation of gap junctions and resumed oocyte maturation in follicles, suggesting that the physical linkages between cumulus cells and oocytes are essential (Kordowitzki et al., 2021; Li et al., 2007). However, it remained unclear how aging is associated with the expression of connexin proteins. At least our data (Figure 2C) and a significant decrease in expression of the cluster of gap junction genes shown by the spatial RNA-seq (Mishina et al., 2021) suggest that an age-dependent reduction in the cumulus cell function diminished the cumulus-oocyte intercellular communications in middle-aged mice.

Our data showed that the impact of aging was more pronounced at fertilization than at post-fertilization. As shown in Figure 3D, the embryo development rate tended to decrease, although this was not significant; average rates were 94.5% and 74.1% in young and middle, respectively (p = 0.569). Transcriptome analysis also failed to show differentially expressed genes with age in the blastocysts (Figure S4A). On the other hand, in humans, previous studies demonstrated that advanced maternal age (above age 35) greatly affected early embryonic development and blastocyst formation (Janny & Menezo, 1996). The proportion of chromosomal aneuploidies, including trisomies, was reported to rise from a 10% incidence at the age of 30 to 20% at the age of 40 in blastocysts in humans (Gruhn et al., 2019; Ntostis et al., 2021). Similarly, the ratio of trisomies and mosaics observed in fetuses was 12.8% from 14-16 months old female mice (Yamamoto et al., 1973); however, the rate of chromosomal abnormalities in oocytes in middle is comparable to those in young (Webster & Schuh, 2017), suggesting that impaired fertilization may contribute to the reduced fertility of middle-aged mice. Although we cannot rule out that embryos obtained from 34- to 38-week-old female mice may have chromosomal defects, the decrease in fertilizing ability should not be overlooked.

The ZP performs various functions during mammalian oogenesis, fertilization, and preimplantation development. Sperm must penetrate the ZP before fertilization. Once fertilization occurs, the mid and inner filaments of ZP fuse to prevent polyspermy (Familiari et al., 1992), and the oocytes secret ovastacin that cleaves ZP2 and hardens the ZP (Burkart et al., 2012). Thus, abnormalities of the ZP cause low fertility or infertility (Familiari et al., 2008; Litscher & Wassarman, 2020). The ZP surface structures have been investigated in various species such as bovine, canine, mouse, and humans (Baez et al., 2019; Nogues et al., 1988; Vanroose et al., 2000). A distinct mesh structure with numerous pores and fenestrations is well conserved across species. Previous studies demonstrated that the successfully fertilized oocytes had ruggedness on the surface structure, while premature oocytes and unfertilized eggs had a smooth structure in human IVF (Familiari et al., 2008; Magerkurth et al., 1999). Our data suggested that the structure of the ZP changed from ruggedness to smoothness, affecting sperm-binding to the ZP and subsequent fertilization in aged mice (Figures 3A, 4A, and C). Consistently with human IVF (Magerkurth et al., 1999), the number of sperm bound to the ZP surface corresponded to the morphology of the ZP (Figure 4C-G). Of note, the destabilization of the ZP structure in aged oocytes by glutathione treatments restored the fertilization rate (Figure 6C).

Our present data show that at middle age there was reduced fertilizing ability of the oocyte due to structural aberrations and dysfunction of the ZP. In addition to the chromosomal abnormalities and subsequent abnormal embryonic development post-fertilization, fertility decline in middle reproductive age is associated with fertilization defects. Thus, improving the follicular environment and restoring fertilizing ability may be a cue for infertility treatment in middle reproductive age.

## Materials and Methods

### Animals

All C57BL/6J mice used in this study were purchased from CLEA Japan (Tokyo, Japan) and Charles River Japan (Yokohama, Japan). Mice were maintained under pathogen-free conditions, under a 12-h light/12-h dark cycle in a temperature-controlled environment of the experimental animal facility at the Institute of Medical Science of the University of Tokyo as well as the Research Institute for Microbial Diseases, Osaka University. All mouse experiments were approved by the Institutional Animal Care and Use Committee of the University of Tokyo (#PA19-10) and the Animal Care and Use Committee of the Research Institute for Microbial Diseases, Osaka University (#Biken-AP-H30-01) and performed in compliance with their guidelines. Young-aged female (young; 10 to 12 weeks old, equivalent to age 20-25 in humans) mice (Dutta & Sengupta, 2016), middle-aged female (middle; 34 to 38 weeks old, equivalent to age 35-40 in humans), and male mice (12- to 15-week-old) were used.

### Smear Test

Vaginal smears from nine young and twenty middle mice were observed every morning (9:00-11:00) and evening (16:00-18:00) continuously for 20 days. Female mice were classified into each stage of the estrous cycle based on the following criteria for the morphology and quantity of cells. The proestrus stage showed nucleated and few cornified epithelial cells, the estrus stage showed many cornified epithelial cells, the metestrus stage showed many leukocytes, and few nucleated epithelial cells, and the diestrus stage showed decreased cell numbers and few leukocytes (Hasegawa et al., 2016).

### Fertility test

Two young- or middle-aged mice were caged with a male mouse two-to-one to observe pregnancy, as modified from a previous work (Hasuwa et al., 2013). Copulation was confirmed by checking for vaginal plugs every morning, and the number of pups was counted after birth. The pregnancy rate was calculated by successful delivery per plug formation.

### Hormone measurement

On the days of estrus and metestrus, 20-50 µL of mouse blood was collected from the tail vein, and serum was obtained after centrifugation. Serum estradiol was analyzed using standard enzyme-linked immunosorbent assay (ELISA). The mouse estradiol kit (KGE014) was purchased from Bio-Techne (Minneapolis, MN) and assays were performed according to the manufacturer’s instruction.

### Observation of ovaries histology and follicular count

For hematoxylin and periodic acid-Schiff (PAS), ovaries were fixed with Boui’’s solution at 4°C overnight. Fixed ovaries were then embedded in paraffin, sectioned by microtome at 5 μm thickness, rehydrated, and treated with 1% periodic acid for 15 min, followed by treatment with Schiff’s reagent (Wako, Osaka, Japan) for 20 min. The sections were stained with hematoxylin prior to imaging. For hematoxylin-eosin (HE) staining, the ovaries were fixed with 4% paraformaldehyde at 4°C overnight. Fixed ovaried were then embedded in paraffin, sectioned by microtome at 5 μm thickness, rehydrated, and treated with hematoxylin for 5 min and eosin for 5 min (Hasuwa et al., 2013). Ovarian sections were observed by using a BZ-X710 microscope (KEYENCE, Osaka, Japan). The number of primordial, primary, secondary, and antral follicles was counted based on the morphology of cell compartments, as described previously (Duncan et al., 2017; Myers et al., 2004).

### Measurement of area of ovarian fibrosis

The area of fibrosis was stained brown in HE-stained sections. The area of fibrosis occupying the whole ovary (%ovarian fibrosis) was measured using ImageJ.

### Oocyte collection, in vitro fertilization, and in vitro culture

Females were superovulated by intraperitoneal injection of pregnant mare’s serum gonadotropin (PMSG; 5 units; ASKA Pharmaceutical, Tokyo, Japan) followed by human chorionic gonadotropin (hCG; 5 units; ASKA Pharmaceutical) 48 h later, as described previously (Miyata et al., 2015). The cumulus-oocyte complexes (COCs) were retrieved from the oviduct 15 h after hCG injection and incubated in HTF medium (ARK Resource, Kumamoto, Japan) until use. In vitro fertilization (IVF) was performed using 100 µL HTF and CARD MEDIUM with the Case 2 method (KYUDO company, Saga, Japan) as described in the instruction manual. Fresh cauda epididymal sperm of males were dispersed in a 300 µL drop of HTF at 37°C for 30 min. Sperm were added to the drop containing the COC at a final concentration of 1.5 × 10^5^ sperm/mL. The oocytes were collected 6 h after IVF and cultured in kSOM (ARK Resource) medium for 5 days with observation every 24-h at 37°C under 5% CO_2_ in air. The numbers of one-cell embryos with pronuclei and unfertilized oocytes were counted and used for estimation of the success rate of fertilization.

### Sperm-Zona binding and Fusion assay

Fresh sperm from the cauda epididymis were dispersed in 200 µL of drops of TYH medium and incubated for 2 h at 37°C under 5% CO_2_ in air to induce capacitation. The final concentration of sperm was adjusted to 1.0 × 10^5^ sperm/mL in 100 µL TYH medium drops for insemination. The sperm-zona pellucida (ZP) binding assay was performed as described previously (Fujihara et al., 2020; Kiyozumi et al., 2020). Briefly, the COCs were treated with hyaluronidase for 5 min to remove cumulus cells and oocytes were washed with TYH medium. The oocytes were then incubated for 2 h with sperm and fixed with 0.25% glutaraldehyde for 30 min. The bound sperm on the ZP were observed with an Olympus IX70 fluorescence microscope and a Keyence BZ-X710 fluorescence microscope. For the fusion assay, the ZP was removed by adding Tyrode (Sigma-Aldrich, Burlington, MA) and the ZP-free oocytes were inseminated with 1.0 × 10^5^ sperm/mL in TYH medium. Twenty minutes after insemination, the ZP-free eggs were gently washed in fresh medium drops and then moved to fresh medium drops, fixed with 0.2% paraformaldehyde, and stained by Hoechst 33342. The eggs were transferred to glass-bottomed chambers to observe sperm nuclei fused with oolemma using a spinning-disk confocal microscope (Olympus) (Miyata et al., 2015).

### Quantitative real-time PCR of isolated cumulus cells

The collection of cumulus cells was performed as described previously (Emori et al., 2020). Briefly, the ovaries from females with 5 IU of PMSG 44-46 h were excised, punctured by using a 26G needle (TERUMO, Tokyo, Japan), and the COCs were gently moved to separate media. The media was composed of bicarbonate-buffered minimal essential medium alpha (MEMα; Thermo Fisher Scientific, Gaithersburg, MD) supplemented with 3 mg/ml of bovine serum albumin (Sigma-Aldrich, St. Louis, MO). Fresh cumulus cells were obtained by removing oocytes from COCs via repeated pipetting using fine-bore pipettes and washed with PBS three times. Total RNA was extracted by using Sepasol^®^-RNA I Super G (nacalai, Tokyo, Japan) according to the manufacturer’s instructions and DNase-treated (Takara Bio Inc., Tokyo, Japan). cDNA was synthesized using SuperScript VILO (Thermo Fisher Scientific) according to the manufacturer’s instructions in a reaction volume of 10 μL. The synthesized cDNA was used for qPCR. qPCR analysis was performed with at least three independent RNA samples using Thunderbird SYBR qPCR Mix (Toyobo, Osaka, Japan) and StepOne system (Thermo Fisher Scientific) for quantification. The fold difference was calculated using the ΔΔCt method (Schmittgen & Livak, 2008). The primers used in this study are shown in Table S2.

### In silico data analysis

GSE159281, a set of single-cell transcriptome data for mouse oocytes and cumulus cells in the antral follicles, was downloaded from the NCBI web site (Mishina et al., 2021). A list of mouse genes encoding gap junctions, which is also reported to be expressed in the cumulus cells, were selected for comparison in this study.

### RNA-seq

Superovulation of females was done using a previous method (Hasegawa et al., 2016). The COCs were recovered from the oviducts 15 h after hCG injection, placed in HTF medium with 1 mg/mL hyaluronidase for 5 min at 37°C under 5% CO_2_. Only 0.5 µL of media containing cumulus cells from each mouse was added to 10 µL lysis buffer (Takara Bio Inc.). The oocytes were then treated with 1 mg/mL collagenase to remove the ZP, as described previously (Yamatoya et al., 2011). For MII oocytes, 20 oocytes from 4 mice (5 oocytes per mouse) were collected in one tube and added to 10 µL lysis buffer. For blastocysts, 20 blastocysts from 4 mice (5 blastocysts per mouse) were collected at 4 days after in vitro fertilization. Total RNAs of each sample were extracted using miRNeasy Micro Kit (QIAGEN). RNA quality was confirmed by using an Agilent 2100 bioanalyzer (Agilent, Santa Clara, CA). The RNA of the cumulus cells extracted from each individual mouse was mixed using equal amounts of RNA. Full-length cDNA was generated using a SMART-Seq HT Kit (Takara Bio) according to the manufacturer’s instructions. An Illumina library was prepared using a NexteraXT DNA Library Preparation Kit (Illumina) according to SMARTer kit instructions. The resulting libraries were sequenced using HiSeq 2500 or NovaSeq 6000 platforms in a single-end mode (Illumina, San Diego, CA, USA). Sequenced reads (approximately 10 million reads) were mapped to the mouse reference genome sequences (mm10) using TopHat version 2.1.1. Normalized FPKM were calculated using Cuffnorm or Cuffdiff version 2.2.1 and each value lower than 0.1 was set to 0.1. Gene Ontology (GO) analysis was performed using Metascape (https://metascape.org/) (Zhou et al., 2019). Hierarchical clustering heatmaps of genes identified in the RNA-seq data was calculated with Ward’s method using Scipy and visualized with Matplotlib. The obtained RNA-seq data have been deposited in the Gene Expression Omnibus database (BioSample accession: SAMD00638922-SAMD00638945).

### Injection of lucifer yellow into COCs

The COCs were collected from the ovaries 44-48 h after PMSG injection and placed in FHM medium until use. The oocyte of each COC was injected with about 10 pL of a 100 mM solution of Lucifer Yellow (L453; Thermo Fisher Scientific), as described previously (El-Hayek et al., 2018). The COCs were incubated for 30 min, then observed using a fluorescence microscope (Keyence). The intensity of GFP in the oocytes and cumulus cells at 0 and 30 min incubation was measured by using ImageJ.

### Glutathione measurement of GV oocytes

The COCs were collected, as described above (*See “Quantitative real-time PCR of isolated cumulus cells”*), and) and placed in TYH medium. The COCs were treated with hyaluronidase (Sigma Type I-S, 150 units/mL) and washed with MEMα medium. Cumulus-free eggs were transferred into 50 µL drops of acidic Tyrode’s solution (Sigma) and pipetted several times until the ZP was dissolved under a stereoscopic microscope (approximately 30 sec). GV oocytes were then washed in PBS with 0.1% PVP three times and stored at –80°C until use. Due to the limitation of detection sensitivity, at least 20 GV oocytes were prepared for one assay. GSH measurements were performed using GSH-Glo™ Glutathione Assay kit (Promega) according to the manufacture’s instruction and SperctraMax L (Molecular Devices, CA).

### Transmission electron microscopy (TEM)

The ovaries were fixed in 2.5% glutaraldehyde + 2% paraformaldehyde in phosphate buffer (0.1 M, pH 7.4) and then fixed with 2.5% glutaraldehyde + 2% paraformaldehyde in phosphate buffer (0.1 M, pH 7.4). The specimens were postfixed in 2% osmium tetroxide (OsO_4_) in phosphate buffer (0.1 M, pH 7.4) at 4°C for 2 h, dehydrated in a graded ethanol series (30%-99%) and then embedded in Epo812 (TAAB). For observation of the ZP filaments, the ovaries and oviducts were pre-soaked in 0.02% saponin and 1.0% ruthenium red in phosphate buffer (0.1 M, pH 7.4) for 30 min, and then fixed with 2.5% glutaraldehyde + 2% paraformaldehyde plus 0.02% saponin and 1.0% ruthenium red in phosphate buffer (0.1 M, pH 7.4) at 4°C overnight (Familiari et al., 2008; Familiari et al., 1989). Ruthenium red is known to act as a staining and stabilizing agent of structural glycoproteins and polyanionic carbohydrates by preventing their dissolution and/or alteration induced by aqueous fixatives (Familiari et al., 1989). The specimens were washed in phosphate buffer (0.1 M, pH 7.4) containing 1.0% RR and 0.02% saponin, postfixed for 1 h in 2.0% OsO_4_ in phosphate (0.1 M, pH 7.4) buffer containing 0.75% ruthenium red and 0.02% saponin and then dehydrated in a graded ethanol series (30%-99%) and then embedded, as described above. Sections (roughly 70 nm thick) were prepared using a Leica EM UC 6 ultramicrotome (Leica, Wetzlar, Germany) and observed using a JEM-1400 Flash transmission electron microscope (JEOL, Tokyo, Japan) at 80 kV (Hirano et al., 2022).

### Measurement of cumulus cell-cell distance

Segmentation of cumulus cells in TEM pictures was performed using Cellpose (https://www.cellpose.org/) (Stringer et al., 2021). The center of cell was calculated, and images were threshold based on the cell shape and then a cell-cell distance was calculated by using ImageJ.

### Scanning electron microscopy (SEM)

SEM analysis was performed, as described previously (Kawano et al., 2010) with some modifications. Briefly, MII oocytes, fertilized eggs, and unfertilized eggs were washed three times with 0.1% PVP/PBS, then with PBS and fixed with 2.0% glutaraldehyde in phosphate buffer (0.1 M, pH 7.4) on poly-l-lysine (PLL; sigma)-coated silicon mounts (G3390 Silicon wafer; EMJapan Co., Ltd., Tokyo, Japan) at room temperature for 1 h. The eggs were treated successively with 1% OsO_4_, 1% Tannic acid, and then 1% OsO_4_ in the above phosphate buffer at 4°C for 10 min and dehydrated with a graded ethanol series (30%-99%). The dehydrated samples were treated with isoamyl acetate and dried in liquid CO_2_ using a critical-point drying apparatus (HCP-2; Hitachi, Tokyo, Japan), transferred onto a stub and coated with osmium coater (HPC-1S; Vacuum Device, Ibaraki, Japan). Images were then acquired under a Zeiss Sigma field-emission SEM (Zeiss Sigma; Carl Zeiss, UK) at a working distance of 5 mm and an accelerating voltage of 5 kV.

### Western blot analysis

Western blotting was conducted, as reported previously (Emori et al., 2020). Following three washes with 0.1% PVP/PBS, twenty MII oocytes were directly mixed with SDS sample buffer (Nacalai) and stored at –80°C until use. Protein lysates were resolved by SDS/polyacrylamide gel electrophoresis (PAGE) and transferred to polyvinylidene fluoride (PVDF) membranes (Bio-Rad, Hercules, CA, USA). After blocking, blots were incubated with anti-ZP2 antibody (1:1000, kindly gifted by Dr. Luca Jovine) at 4°C for overnight and then incubated with secondary antibodies conjugated with horseradish-peroxidase (HRP) or conjugated donkey anti-rat IgG [1:2000, 712-035-153; Jackson ImmunoResearch, West Grove, PA, USA]. GAPDH was used as an internal control (1:1000, Direct-Blot™ HRP anti-GAPDH antibody, 607903; Biolegend, San Diego, CA, USA). Signals were visualized using Luminata Forte (Merck Millipore, Tokyo, Japan) and the ChemiDoc Imaging System (Bio-Rad).

### Mass spectrometry and data analysis

Isolation of the ZP was performed by using a piezo-manipulator (Yamagata et al., 2002). Briefly, the COC was retrieved from the oviduct 15 h after hCG injection, as described above (See “*Oocyte collection, in vitro fertilization, and in vitro culture”*), and placed in HEPES-buffered KSOM (FHM) at room temperature. The COCs were treated with hyaluronidase and oocytes were washed with FHM medium. The ZP was drilled by a piezo-manipulator PMAS-CT150 (Prime Tech LTD., Ibaraki, Japan) in FHM with 0.25 M sucrose and then separated from the oocyte. Five the ZP collected from each mouse, in three replicates, were subjected to MS analysis. The ZP samples were purified using methanol-chloroform precipitation and then dissolved in 0.1% Rapigest solution (Waters, Milford, MA, USA). The ZP proteins were reduced with 10 mM dithiothreitol (DTT), followed by alkylation with 55 mM iodoacetamide, and digested by treatment with trypsin and purified with a C18 tip (GL-Science, Tokyo, Japan). The resultant peptides were subjected to nanocapillary reversed-phase LC-MS/MS analysis using a C18 column (12 cm x 75 um, 1.9µm, Nikkyo technos, Tokyo, Japan) on a nanoLC system (Bruker Daltoniks, Bremen, Germany) connected to a tims TOF Pro mass spectrometer (Bruker Daltoniks) and a modified nano-electrospray ion source (CaptiveSpray; Bruker Daltoniks). The mobile phase consisted of water containing 0.1% formic acid (solvent A) and acetonitrile containing 0.1% formic acid (solvent B). Linear gradient elution was carried out from 2% to 35% solvent B for 20 min at a flow rate of 250 nL/min. The ion spray voltage was set at 1.6 kV in the positive ion mode. Ions were collected in the trapped ion mobility spectrometry (TIMS) device over 100 ms and MS and MS/MS data were acquired over an *m/z* range of 100-2,000. During the collection of MS/MS data, the TIMS cycle was adjusted to 0.53 s and included 1 MS plus 4 parallel accumulation serial fragmentation (PASEF)-MS/MS scans, each containing on average 12 MS/MS spectra (>100 Hz) (Meier et al., 2015; Meier et al., 2018), and nitrogen gas was used as collision gas. The resulting data was processed using Data Analysis version 5.2 (Bruker Daltoniks), and proteins were identified using MASCOT version 2.7.0 (Matrix Science, London, UK) against the Uniprot mouse database (88,017 sequences; 35,135,138 residues). Protease specificity was set for trypsin (C-term, KR; Restrict, P; Independent, no; Semispecific, no; two missed and/or nonspecific cleavages permitted). Variable modifications considered were N-terminal Gln to pyro-Glu, and oxidation of methionine. The mass tolerance for precursor ions was ±15 ppm. The mass tolerance for fragment ions was ±0.05 Da. The threshold score/expectation value for accepting individual spectra was p < 0.05. Quantitative value and fold exchange were calculated by Scaffold5 version 5.1.2 (Proteome Software, Portland, OR, USA; Protein Threshold: 99%; Min # Peptides: 2; Peptide Threshold: 95%) for MS/MS-based proteomic studies (Searle, 2010). The quantitative values (Normalized Total Spectra) are available in Table S1.

### Analysis of the fractal dimension

The fractal analysis allows the complexity of shapes and pattern recognition to be quantified as fractal dimensions (Mandelbrot, 1967). Fractal dimensions of the ZP surface structure and cross-images were analyzed using a custom-written MATLAB-based algorithm based on the library (Dennis & Dessipris, 1989): https://jp.mathworks.com/matlabcentral/fileexchange/71774-create-measure-characterize-visualize-1d-2d-3d-fractals. Note that this library is only applicable to the calculation of fractal dimensions. The code for fractal analysis generated during this study are available at the GitHub: https://github.com/YuIshikawaYamauchi/Ishikawa_et_al_2023. The fractal dimension ranged from 1 to 3, with bin edges in increments of 0.2. In addition to the fractal dimension, our custom algorithm read in images, calculated image features, including the number of holes and the edge intensity, and performed linear multiple regression training and prediction. The imported original images were first subjected to smoothing image processing (https://jp.mathworks.com/help/images/ref/imbilatfilt.html). The fractal dimensions define the median of all pixels as the fractal dimension representative of the image after calculating the fractal dimension per pixel. The calculation of the fractal dimension per pixel was performed in the following steps: all pixels D1 (difference from neighboring pixels) and D2 (difference from pixels epsilon away) were calculated. For D1 and D2, the average values Dm1 and Dm2 of the neighborhood blocks of the window width were calculated, and the fractal dimension was calculated using the following formula.

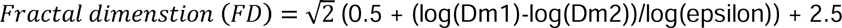

In the area histogram of holes, the image was converted to a binary image, and the number of holes per area was represented by histograms. Based on the results of the validation, the optical histogram bin edges were set to 0, 50, 100, 150, 1000, and 1000000. In the calculation of edge intensity, the number of edges per intensity was represented by histograms. The edges of the histogram bins were 0, 20, 40, 60, and 100, but the weights were all set to zero in this study, as the edge intensities were not of much importance when verified. The reasons as follows; there were 19 explanatory variables including 10 bins for the histogram of fractal dimension, 5 bins for the histogram of the number of holes, and 4 bins for the edge strength. Among these 19 variables, effective variables for linear regression were selected, and the weight of the unselected variables was fixed to zero. Specifically, we performed the following processing, and the resulting setE_best became the selected explanatory variables, while the weight of the unselected variables became zero.

1. Upon selection of all variables, the set of selected variables was setE. The mean absolute error (MAE) of the Leave One Out (LOO) cross-validation was then determined and initialized as follows.

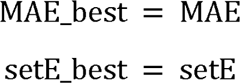
2. LOO cross-validation was performed by excluding one variable from the variables stored in setE, and the best MAE with the highest accuracy was set as MAE_best_tmp.
3. setE was updated as follows: the set of explanatory variables when setE = MAE_best_tmp (the one with the best accuracy among the ones obtained by excluding one variable from setE).
4. If MAE_best > MAE_best_tmp, the following updates are performed:

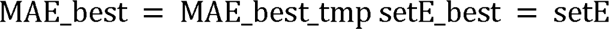
5. The process ended when the number of variables in setE became one. Otherwise, the process returned to step 2.”

As learning YAF and MAF using linear multiple regression, the following variables were trained. Explanatory variable X is a standardized histogram of the three types of features, fractal dimension histogram, vacancy area histogram, and edge intensity histogram. The objective variable Y is YAF set to 0 and MAF set to 1. The linear multiple regression equation was as follows.

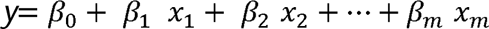

Ideally, the estimate y should be 0 or 1, but in practice, y can take values below 0 or above 1, so it is transformed into a likelihood, p, by the following sigmoidal function.

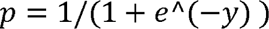

As mentioned above, we used Leave One Out (LOO) cross-validation to test the accuracy of our machine-learning models on unknown data not used for training. The LOO cross-validation test performs the following tests N times: 1. create a training model with N-1 teacher data by excluding one piece of data from N teacher data; 2. input the excluded one piece of data into the training model to predict the objective variable; and 3. perform the same test N times to estimate all the data. Since all these estimation results are not included in building the learning model, the prediction accuracy of the LOO cross-test can be regarded as the prediction accuracy for the unknown data.

### Statistical analysis

Statistical analysis was performed with the GraphPad Prism 9 software. Data are represented as mean ± SEM of three or more biological replicates. Statistical tests are detailed in each Figure legend. Differences were considered significant at \**p* < 0.05, ** *p* < 0.01, and *** *p* < 0.001. Error bars represent SEM. Figure legends indicate the number of n values and the number of experiments for each analysis.

## Acknowledgments

The authors thank Dr. Yasuhiro Yamada for valuable advice and discussions, Miyako Baba and Mio Kikuchi for technical assistance. Drs. Natsuko Kawano, Masanori Nasu, and

Fusako Mitsuhashi for technical assistance for SEM. Drs. Yoshihiro Kawaoka and Michiko Ujie for usage of SEM. Dr. Julio M. Castaneda and Dr. Daiji Kiyozumi for critical reading of the manuscript. This work was supported by the Ministry of Education, Culture, Sports, Science and Technology (MEXT)/Japan Society for the Promotion of Science (JSPS) KAKENHI grants (JP19J01089 to Y.IY., and JP19H05750 and JP21H05033 to M.I.), Japan Agency for Medical Research and Development (AMED) grant JP21gm5010001 to M.I., the Core Research for Evolutional Science and Technology (CREST)/Japan Science and Technology Agency (JST) grant (JPMJCR21N1) to M.I.

## Author Contributions

Y.IY. and M.I. conceived the study and wrote the manuscript together. Y.IY., C.E., K.K., and T.E. performed the research and analyzed the data. D.M. and H.M. performed bulk RNA-seq analysis. Y.W. and H.S. assisted and performed TEM and SEM. T.N. developed the algorithm for fractal analysis. A.N. performed MS analysis. M.O. assisted and advised on the research.

