## Supplemental files for "Age-associated aberrations of cumulus-oocyte interaction and microfilamentous structure in the zona pellucida decline female fertility"

### Supplemental Figure 1

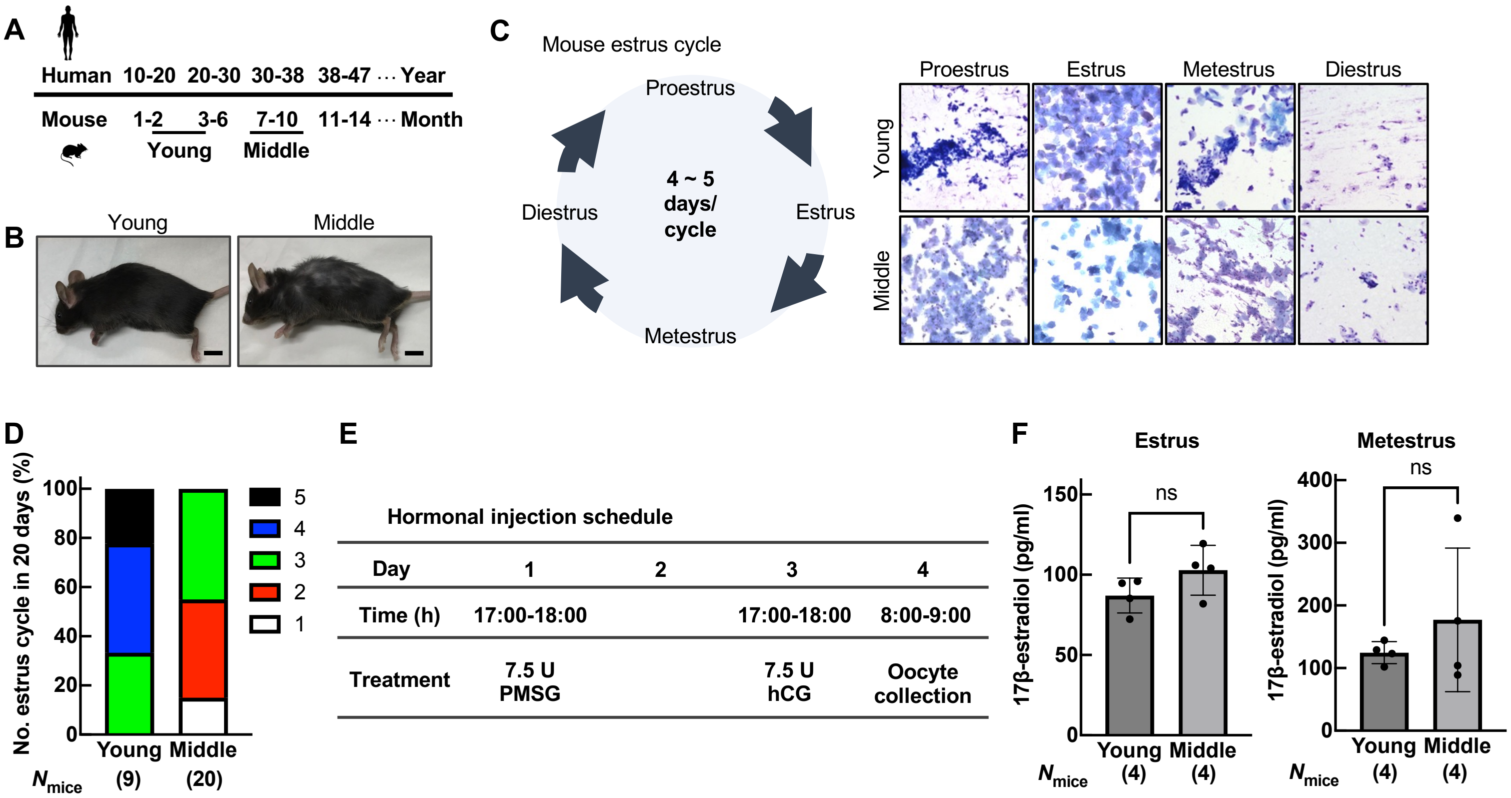


**Figure S1. Behavior of estrus cycles and hormonal status in middle-aged mice.**
(A) Analogical age relationship between human and mice (Dutta and Sengupta, 2016). (B) Appearance of young-aged female (10- to 12-week-old; young) and middle-aged female (34- to 38-week-old; middle) mice. Middle-aged mice have noticeable hair loss on their body hair in appearance. Scale bars = 10 mm. (C) The mouse estrus cycle. Giemsa staining of smear cells. Mouse estrus cycle was divided into four stages including the proestrus, estrus, metestrus, and diestrus. Based on the morphology of the cells, each stage were defined. (D) The number of estrus cycles in 20 consecutive days. (E) Time schedule of superovulation to mice and collection of the oocytes. (F) Endogenous 17β-estradiol levels at the estrus and metestrus. There was no significant difference between young and middle mice. ns, not significant ($p>0.05$; unpaired Student's t-test). Data are mean ± SEM.

### Supplemental Figure 2

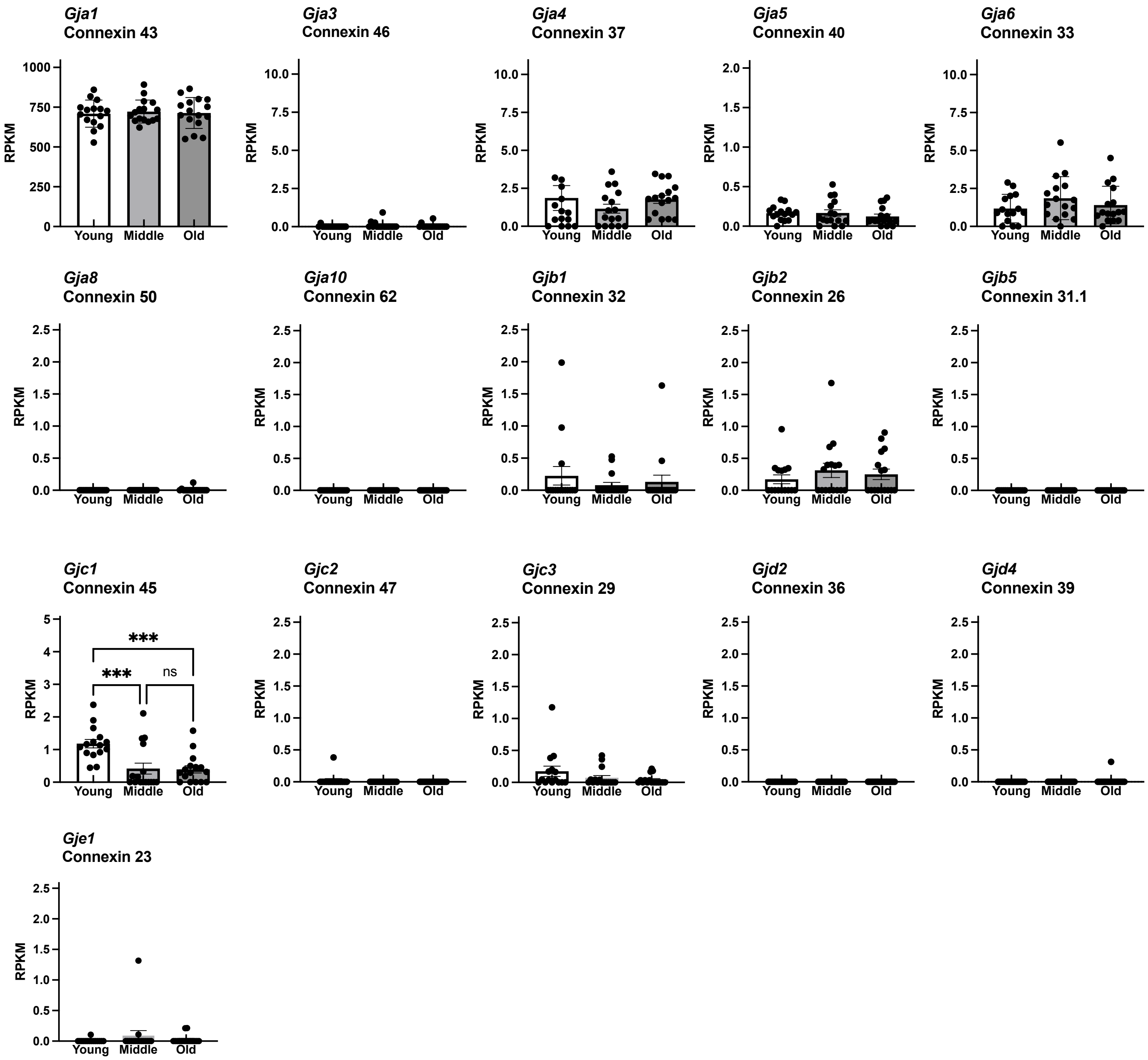


**Figure S2. Expression of gap junction genes among young, middle, and old mice.**
scRNA-seq of cumulus cells in the antral follicles from 2 months (Young), 9 months (Middle), and 14 months (Old) mice. scRNA-seq data were obtained from GSE159281. *** $p<0.001$; ns, not significant ($p>0.05$, Two-way ANOVA with Tukey's multiple comparisons test). Data are mean ± SEM.

### Supplemental Figure 3

**A**

**Hormonal injection schedule**

| Day | 1 | 2 | 3 | 4 | 5 | 6 | 7 |
|---|---|---|---|---|---|---|---|
| Time (h) | 17:00-18:00 | 17:00-18:00 | 17:00-18:00 | 17:00-18:00 | | 17:00-18:00 | 8:00-9:00 |
| Treatment | P4 | P4 | AIS | AIS | | hCG | Oocyte collection |

**B**

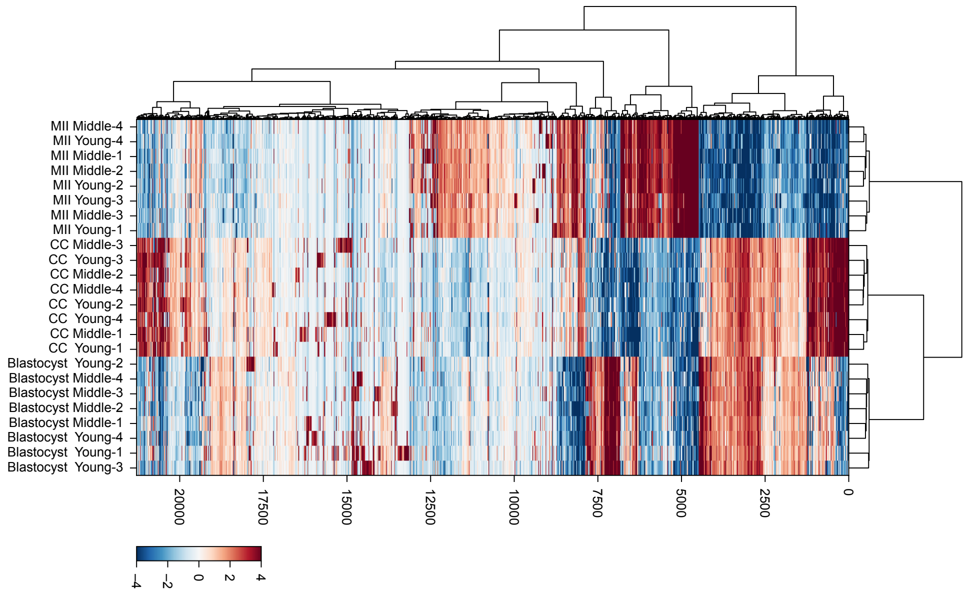


**C**

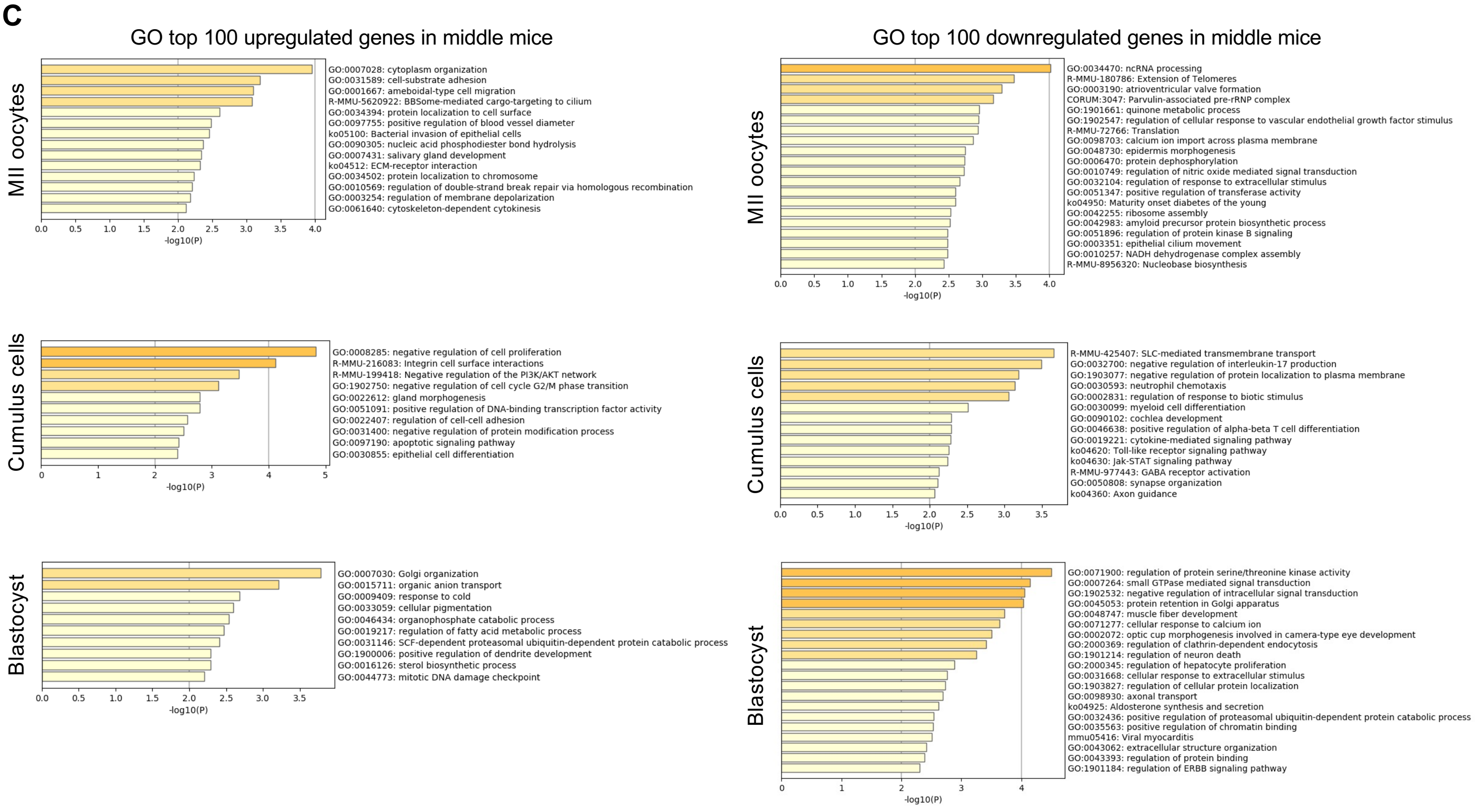


**Figure S3 Transcriptome analysis of MII oocytes and cumulus cells at ovulation and blastocyst after IVF.**
(A) Hormonal injection for collecting a large number of MII oocytes (Hasegawa et al., 2016). Progesterone, P4; Anti-inhibin serum, AIS. (B) Bulk RNA-seq analysis of MII oocytes and cumulus cells at ovulation and blastocysts after IVF from young and middle mice. Heatmap showing changes in expression levels of upregulated and downregulated genes of MII oocytes, cumulus cells, and blastocysts (Fold change>2, *p*<0.05 [FDR-adjusted Wald Test]). (C) Gene ontology (GO) term enrichments analysis of top 100 upregulated and downregulated genes in MII oocytes, cumulus cells, and blastocysts. Metascape (https://metascape.org/) was used for the analysis.

### Supplemental Figure 4

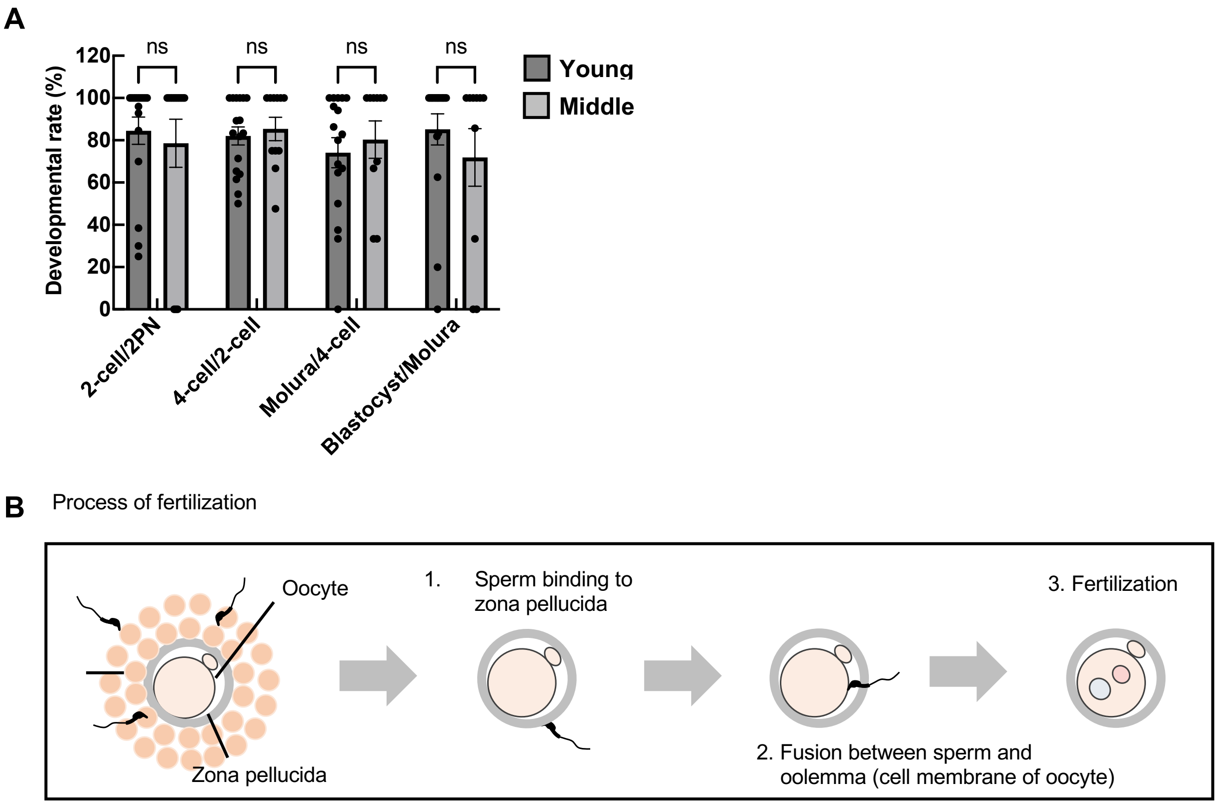


**Figure S4. Developmental rate and process of fertilization.**
(A) Developmental rate from 2PN to blastocyst. ns, not significant ($p$>0.05, Two-way ANOVA with Tukey's multiple comparisons test). (B) Process of fertilization. Data are mean ± SEM.

### Supplemental Figure 5

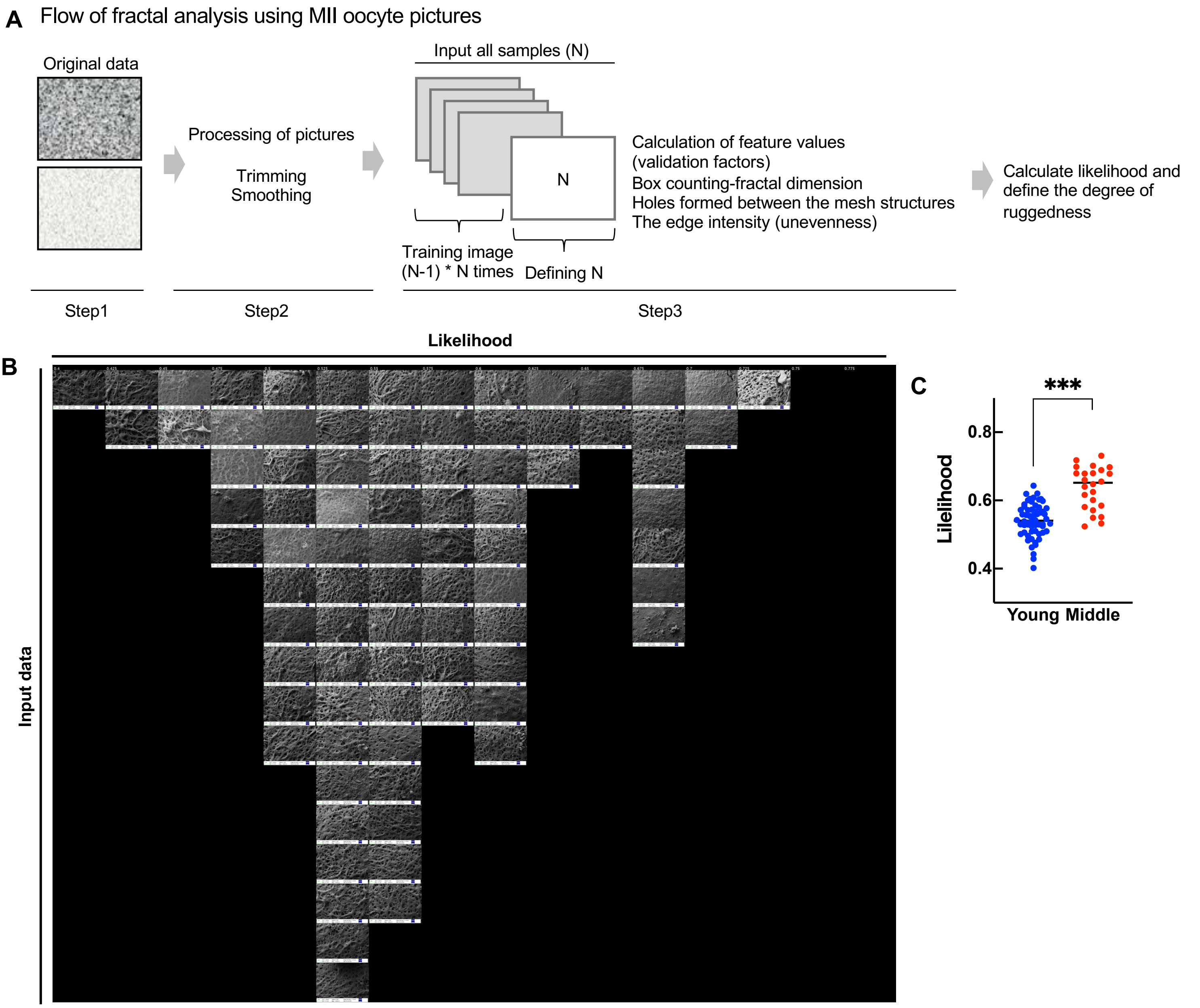


**Figure S5. Computational analysis of the surface ruggedness in the ZP.**
(A) Schematic diagram of the the surface ruggedness automatic discrimination tool, the fractal analysis tool. In this study, we used a method to discriminate the degree of ruggedness of one image by training with N-1 images using 83 samples (leave-one-out cross-validation/LOOCV). The likelihood gives a numerical value for the degree of ruggedness. In the fractal analysis of this tool, the calculated numerical value indicates the ruggedness. If the image is irregular, the numerical value is small, and if it is regular, the numerical value is large. This indicates that MII oocytes of middle-aged mice are discriminated as having a regular surface and small ruggedness as an image. (B) All images used for fractal analysis, which refers to step 3 of (A). (C) Each likelihood of yound and middle was calculated using the fractal analysis tool. *** $p<0.001$, Tukey-Kramer test.

### Supplemental Figure 6

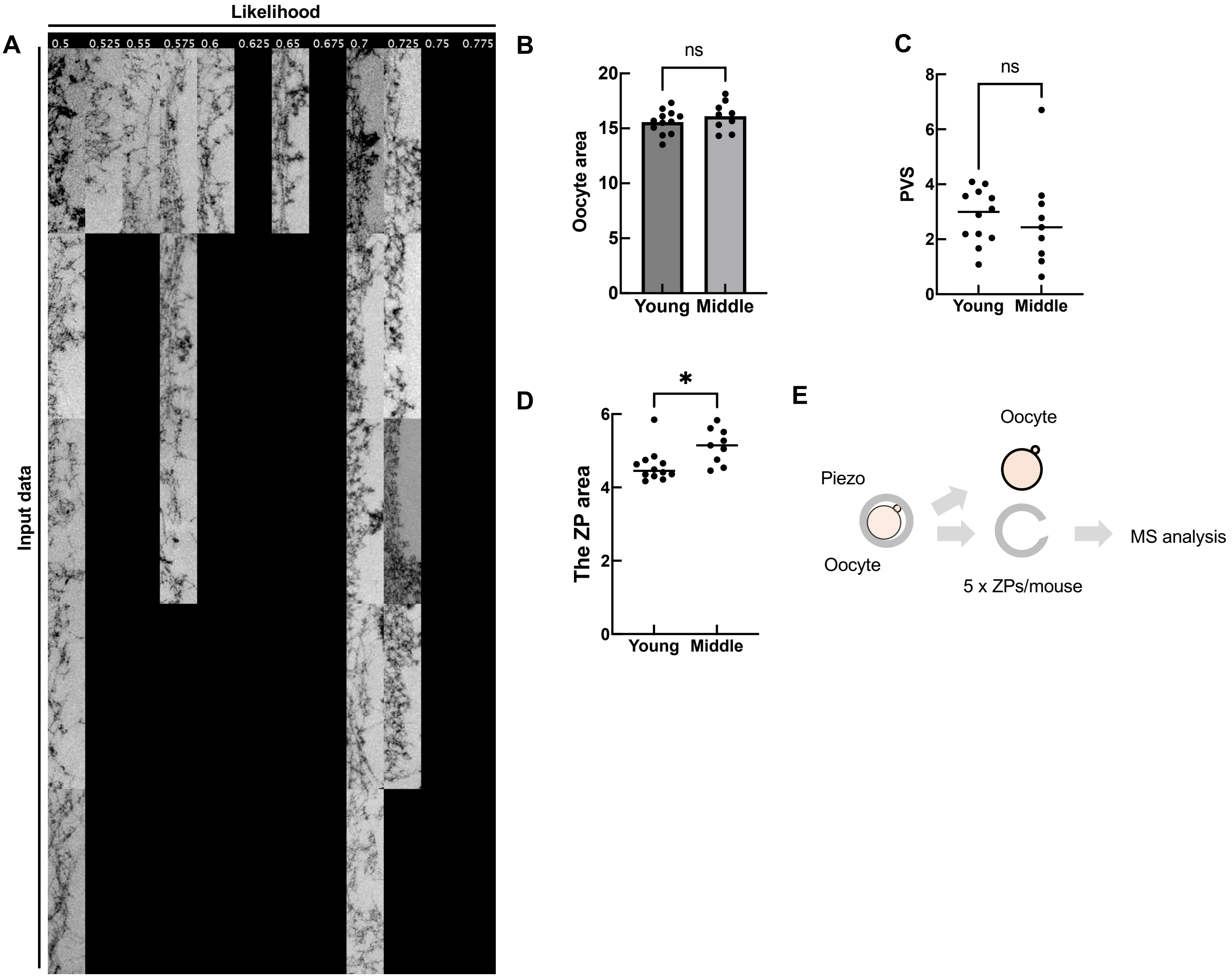


**Figure S6. Characteristics of the ZP in middle-aged mice.**

(A) All images of TEM used for fractal analysis. (B) The area (pixel) of oocyte including outer layer of the ZP. ns, not significant ($p$>0.05; Tukey-Kramer test). (C) Calculated area (pixel) of perivitelline space in the oocyte. ns, not significant ($p$>0.05; Tukey-Kramer test). D) Total area (pixel) of the ZP. *$p$<0.05; Tukey-Kramer test. (E) Schematic diagram for MS analysis of MII oocytes. a piezo-manipulator was used for drilling the ZP. Five the ZP per mouse was collected and applied to MS analysis.

### Supplemental Table 1

#### Table 1. MS analysis of the ZP

| Identified Proteins (keratins excluded) | Accession Number | Alternate ID | Molecular Weight | Quantitative Value (Normalized Total Spectra) | | | | | | |
|---|---|---|---|---|---|---|---|---|---|---|
| | | | | Young | | | Middle | | | |
| | | | | exp_01 | exp_02 | exp_04 | exp_01 | exp_02 | exp_03 | |
| Serum albumin | Q546G4 | Alb | 69 kDa | 38.852 | 83.397 | 14.228 | 13.787 | 99.527 | 4.9645 | Serum albumin OS=Mus musculus OX=10090 GN=Alb PE=1 SV=1 |
| Zona pellucida sperm-binding protein 2 | P20239 | Zp2 | 80 kDa | 16.109 | 18.741 | 17.073 | 22.059 | 25.877 | 29.787 | Zona pellucida sperm-binding protein 2 OS=Mus musculus OX=10090 GN=Zp2 PE=1 SV=1 |
| Zona pellucida sperm-binding protein 3 | P10761 | Zp3 | 46 kDa | 12.319 | 19.678 | 32.724 | 24.816 | 20.901 | 22.34 | Zona pellucida sperm-binding protein 3 OS=Mus musculus OX=10090 GN=Zp3 PE=1 SV=4 |
| Zona pellucida sperm-binding protein 1 | Q62005 | Zp1 | 69 kDa | 6.6332 | 13.119 | 14.228 | 14.706 | 15.924 | 22.34 | Zona pellucida sperm-binding protein 1 OS=Mus musculus OX=10090 GN=Zp1 PE=1 SV=1 |
| Predicted pseudogene 5478 | A0A2R8VHP3 | Gm5478 | 58 kDa | 16.109 | 9.3705 | 4.2683 | 11.03 | 9.9527 | 7.4468 | Predicted pseudogene 5478 OS=Mus musculus OX=10090 GN=Gm5478 PE=1 SV=1 |
| Annexin A2 OS=Mus musculus | P07356 | Anxa2 | 39 kDa | 1.8952 | 27.174 | 1.4228 | 2.7574 | 0 | 0 | Annexin A2 OS=Mus musculus OX=10090 GN=Anxa2 PE=1 SV=2 |
| Heat shock 70 kDa protein 1B | P17879 | Hspa1b | 70 kDa | 4.738 | 7.4964 | 0 | 5.5148 | 1.9905 | 0 | Heat shock 70 kDa protein 1B OS=Mus musculus OX=10090 GN=Hspa1b PE=1 SV=3 |
| Protease, serine, 3 | B9EJ35 | Prss3 | 26 kDa | 1.8952 | 4.6852 | 7.1138 | 2.7574 | 2.9858 | 9.9291 | Protease, serine, 3 OS=Mus musculus OX=10090 GN=Prss3 PE=2 SV=1 |
| L-lactate dehydrogenase C chain | P00342 | Ldhc | 36 kDa | 1.8952 | 0.93705 | 4.2683 | 2.7574 | 1.9905 | 0 | L-lactate dehydrogenase C chain OS=Mus musculus OX=10090 GN=Ldhc PE=1 SV=2 |
| Desmoplakin OS=Mus musculus | E9Q557 | Dsp | 333 kDa | 4.738 | 1.8741 | 0 | 3.6765 | 1.9905 | 0 | Desmoplakin OS=Mus musculus OX=10090 GN=Dsp PE=1 SV=1 |
| Oviduct-specific glycoprotein | Q62010 | Ovgp1 | 79 kDa | 0 | 0 | 2.8455 | 0.91913 | 0 | 0 | Oviduct-specific glycoprotein OS=Mus musculus OX=10090 GN=Ovgp1 PE=2 SV=1 |
| Clusterin OS=Mus musculus | Q06890 | Clu | 52 kDa | 0 | 0.93705 | 1.4228 | 1.8383 | 1.9905 | 0 | Clusterin OS=Mus musculus OX=10090 GN=Clu PE=1 SV=1 |
| Lactadherin OS=Mus musculus | P21956 | Mfge8 | 51 kDa | 0.94761 | 0.93705 | 2.8455 | 5.5148 | 1.9905 | 4.9645 | Lactadherin OS=Mus musculus OX=10090 GN=Mfge8 PE=1 SV=3 |
| Actin, beta | B2RRX1 | Actb | 42 kDa | 0 | 0 | 0 | 2.7574 | 0.99527 | 0 | Actin, beta OS=Mus musculus OX=10090 GN=Actb PE=2 SV=1 |
| Succinyl-CoA:3-ketoacid coenzyme A transferase 2B, mitochondrial | Q9ESL0 | Oxct2b | 57 kDa | 0.94761 | 0 | 1.4228 | 1.8383 | 2.9858 | 0 | Succinyl-CoA:3-ketoacid coenzyme A transferase 2B, mitochondrial OS=Mus musculus OX=10090 GN=Oxct2b PE=2 SV=1 |
| A-kinase anchor protein 4 | Q60662 | Akap4 | 94 kDa | 0 | 0 | 0 | 0 | 0 | 0 | A-kinase anchor protein 4 OS=Mus musculus OX=10090 GN=Akap4 PE=1 SV=1 |
| Junction plakoglobin | Q02257 | Jup | 82 kDa | 1.8952 | 0 | 0 | 3.6765 | 2.9858 | 0 | Junction plakoglobin OS=Mus musculus OX=10090 GN=Jup PE=1 SV=3 |
| Histone H4 | B2RTM0 | Hist2h4 | 11 kDa | 0.94761 | 0 | 0 | 5.5148 | 0.99527 | 0 | Histone H4 OS=Mus musculus OX=10090 GN=Hist2h4 PE=1 SV=1 |
| 78 kDa glucose-regulated protein | Q3TWF2 | Hspa5 | 72 kDa | 0 | 5.6223 | 0 | 0 | 0 | 0 | 78 kDa glucose-regulated protein OS=Mus musculus OX=10090 GN=Hspa5 PE=2 SV=1 |
| Regucalcin | Q64374 | Rgn | 33 kDa | 0 | 3.7482 | 0 | 0 | 0 | 0 | Regucalcin OS=Mus musculus OX=10090 GN=Rgn PE=1 SV=1 |
| Catalase | Q3TVZ1 | Cat | 60 kDa | 0 | 0 | 0 | 1.8383 | 0 | 0 | Catalase OS=Mus musculus OX=10090 GN=Cat PE=2 SV=1 |
| Transaldolase | Q93092 | Taldo1 | 37 kDa | 0 | 10.308 | 0 | 0 | 0 | 0 | Transaldolase OS=Mus musculus OX=10090 GN=Taldo1 PE=1 SV=2 |

### Supplemental Table 2

**Table 2. Primer sets for qPCR of cumulus cells in the antral follicles.**

| Gene name | | Primer sequence |
|---|---|---|
| Cyp19a1_Fw | Fw | ATGTTCTTGGAAATGCTGAACCC |
| Cyp19a1_Rv | Rv | AGGACCTGGTATTGAAGACGAG |
| DDX4_Fw | Fw | GCTTCATCAGATATTGGCGAGT |
| DDX4_Rv | Rv | GCTTGGAAAACCCTCTGCTT |
| Cx43_gap junction protein, alpha 1_Fw | Fw | ACAGCGGTTGAGTCAGCTTG |
| Cx43_gap junction protein, alpha 1_Rv | Rv | GAGAGATGGGGAAGGACTTGT |
| Cx37_gap junction protein, alpha 4_Fw | Fw | CCCACATCCGATACTGGGTG |
| Cx37_gap junction protein, alpha 4_Rv | Rv | CGAAGACGACCGTCCTCTG |
| Cx45_gap junction protein, gamma 1_Fw | Fw | AGATCCACAACCATTCGACATTT |
| Cx45_gap junction protein, gamma 1_Rv | Rv | TCCCAGGTACATCACAGAGGG |
| Gapdh_Fw | Fw | ATGAATACGGCTACAGCAACAGG |
| Gapdh_Rv | Rv | CTCTTGCTCAGTGTCCTTGCTG |
